# *Acat1* gene KO restores TGN cholesterol deficiency in mutant NPC1 cells and expands mutant *Npc1* mouse lifespan

**DOI:** 10.1101/2020.08.07.241471

**Authors:** Maximillian A. Rogers, Catherine C.Y. Chang, Robert A. Maue, Elaina M. Melton, Andrew A. Peden, William S. Garver, Mitchell M. Huang, Peter W. Schroen, Ta-Yuan Chang

## Abstract

Niemann-Pick type C (NPC) is a neurological disorder with no cure. NPC proteins deliver cholesterol from endosomes to other compartments including trans-Golgi network (TGN) and endoplasmic reticulum (ER). Acyl-coenzyme A:cholesterol acyltransferase 1 (ACAT1) is a resident ER enzyme that converts cholesterol to cholesteryl esters for storage. Here, we report the surprising finding that in a mutant *Npc1* mice, *Acat1*-deficiency delayed the onset of weight loss and declining motor skill, prolonged lifespan, delayed Purkinje neuron death, and improved hepatosplenic pathology. Furthermore, syntaxin 6, a cholesterol-binding t-SNARE normally localized to TGN, is mislocalized in mutant NPC cells. However, upon ACAT1 inhibition this mislocalization is corrected, and increase the level of a few proteins further downstream. Our results imply that ACAT1 inhibition diverts a cholesterol storage pool in a way that replenished the low cholesterol level in NPC-deficient TGN. Taking together, we identify ACAT1 inhibition as a potential therapeutic target for NPC treatment.

## Introduction

Niemann-Pick disease type C (NPC) is a genetically recessive neurodegenerative disease caused by mutations in *Npc1* (1), (2) or in *Npc2* (3). Loss of in NPC1 or NPC2 function results in the accumulation of cholesterol (4) as well as various sphingolipid species (5), mainly within late endosomes/lysosomes (LE/LYS). NPC disease shares many similar attributes with Alzheimer’s disease, and is colloquially referred to as juvenile Alzheimer’s disease. As such any mechanistic and therapeutic findings in NPC disease may have broad application to other neurodegenerative diseases. The lipid accumulation seen in NPC occurs in all tissues, and results in neurodegeneration as well as malfunctions in liver and lung. In brain, the most extensive cell death occurs in cerebellum, with preferential loss of Purkinje neurons. In terms of potential therapies, migluostat, a glycosphingolipid synthesis inhibitor, has demonstrated significant efficacy (6). However, migluostat is not FDA-approved as an NPC therapy. Intrathecal delivery of 2-hydroxypropyl-β-cyclodextrin, a water-soluble molecule that binds cholesterol, reduces neurological disease progression (7), (8), but the clinical benefit of cyclodextrin has yet to be clearly demonstrated in ameliorating the disease pathology in NPC1 patients. Given the current lack of approved NPC treatments, there remains a critical need to develop therapeutic approaches for the treatment of NPC disease.

Cholesterol is an essential lipid molecule needed for cell growth and function. Cells acquire cholesterol through exogenous sources as well as through *de novo* biosynthesis. Exogenous cholesterol enters cells mainly through receptor-mediated endocytosis. Subsequently, it is distributed to various membrane compartments for utilization, feed-back regulation of cholesterol metabolism, and storage as cytoplasmic cholesteryl ester lipid droplets [Reviewed in (9)]. Distribution of cholesterol from the LE/LYS to other membrane compartments requires NPC1 and NPC2. Both NPC1 and NPC2 bind to cholesterol (10), (11), (12), (13). These two proteins work in concert to export cholesterol from LE to other membrane organelles [reviewed in (14)]. In cells with NPC mutations, buildup of cholesterol and other lipids occurs within LE/LYS, while leaving other membrane compartments, including plasma membrane (PM) (15), (16), endoplasmic reticulum (ER) (17), (18), (19), (20), peroxisomes (21), and trans-Golgi network (TGN) (22), (23) relatively deficient in cholesterol. In mutant NPC cells, abnormal membrane cholesterol distribution cause malfunctions in LE/LYS (24), and in other membrane organelles (23), (25). Cholesterol overload in LE/LYS also causes cholesterol accumulation in the inner membranes of mitochondria (26), (27), (28), (29).

In addition to receiving cholesterol from exogenous uptake, cells produce cholesterol from *de novo* biosynthesis. Upon synthesis at the ER, sterols quickly move to the PM within a few minutes, via mechanisms independent of NPC1 (30). Cholesterol from both endogenous and exogenous sources traverse among various membrane compartments [reviewed in (31), (32)]. To prevent overaccumulation of free cholesterol in cells, which would result in cellular toxicity [reviewed in (33), acyl-coenzyme A:cholesterol acyltransferase 1 (ACAT1) (also called sterol O-acyltransferase 1 [SOAT1]) (34), converts a certain portion of cellular cholesterol to cholesterol esters. Furthermore, ATP binding cassette transporter A1 (ABCA1) [reviewed in (35)]. removes excess cholesterol through a lipid efflux process.

ACAT1 is a membrane protein residing at the ER (36); in addition, a certain portion of ACAT1 is found close to other cellular organelles including PM (37), recycling endosomes, (38), and TGN (39). In mutant NPC cells, the absence of functional NPC1 or NPC2 considerably slows delivery of cholesterol from LE/LYS to ER; however, a significant amount of cholesterol can translocate from the PM to the ER for esterification in an NPC-independent manner (16, 40-43), (44, 45). Here, we hypothesize that in mutant NPC cells, ACAT1 inhibition causes the NPC-independent cholesterol pool arriving at ER to build up. Once built-up, this cholesterol pool moves away from ER to other subcellular membrane compartments, including TGN, to fulfill their needs for cholesterol. To test this hypothesis, we adopted a transgenic mouse model-based approach.

A mutant mouse model for NPC disease (*Npc1^nmf^* mouse) was discovered and characterized by Maue *et al*. (46). This *Npc1^nmf^* mouse model has a C57BL/6J genetic background, and a single D1005G-*Npc1* mutation located within the cysteine-rich luminal loop of the NPC1 protein; which is comparable to mutations that commonly occur in human *Npc1* patients. Homozygous *Npc1^nmf/nmf^* mice begin to die by 90 days (almost 13 weeks) after birth; and exhibit a phenotype that mimics the late onset, slowly progressing form of NPC disease. In contrast, heterozygous *Npc1^nmf/+^* mice seem to exhibit a generally normal mouse phenotype and are fertile. In the studies presented here, we bred heterozygous *Npc1^nmf/+^* mice with global *Acat1^-/-^* (*A1^-/-^)*-deficient mice (47), which also have a C57BL/6J genetic background, to produce *Npc1^nmf/nmf^*:*A1*^+/+^ and *Npc1^nmf/nmf^*:*A1^-/-^* mice. We then performed paired studies *in vivo* by using sex and age matched *Npc1^nmf/nmf^*:A1^+/+^ (*Npc1^nmf^*) and *Npc1^nmf/nmf^*:*A1^-/-^* mice (*Npc1^nmf^*:*A1^-/-^)*. We used littermates from *Npc1^+/+^:A1^+/+^* mice (WT) and *Npc1^+/+^*:*A1*^-/-^ mice (*A1^-/-^*) produced from the same breeding experiments as non-diseased controls. In addition, we isolated embryonic fibroblast cells from *Npc1^nmf^, Npc1^nmf^:A1^-/-^*, WT, and *A1^-/-^* mice to perform paired studies *in vitro* in primary cell culture. Furthermore, to evaluate the relevance of our findings in the context of human disease, we monitored the effect of a small molecule ACAT1-specific inhibitor K604 on human fibroblast (Hfs) cells isolated from several patients with NPC disease as well as on a human fibroblast cells isolated from a patient with a related lysosomal storage disease Niemann-Pick type A (NPA). The results of both these *in vivo* and *in vitro* approaches are reported here.

## Results

### *Acat1* gene deficiency (*A1*^-/-^) increased life span and reduced weight loss in *Npc1^nmf^* mice

To evaluate effects of *A1***^-/-^** on lifespan of homozygous *Npc1^nmf^* mice, mice with four genotypes (WT, *A1^-/-^, Npc1^nmf^*, and *Npc1^nmf^*:*A1^-/-^*) were fed a regular chow diet and their lifespans were assessed. The age of death for *Npc1^nmf^* mice with or without *A1* were determined as the point where mice could no longer take in food or water, as described previously (46). Results (**Fig. 1A**) show that the median survival for *Npc1^nmf^* and *Npc1^nmf^*:*A1*^-/-^ mice is 113 days and 138 days respectively, with the mean survival being 102 days and 137 days, respectively.

**Fig. 1.**
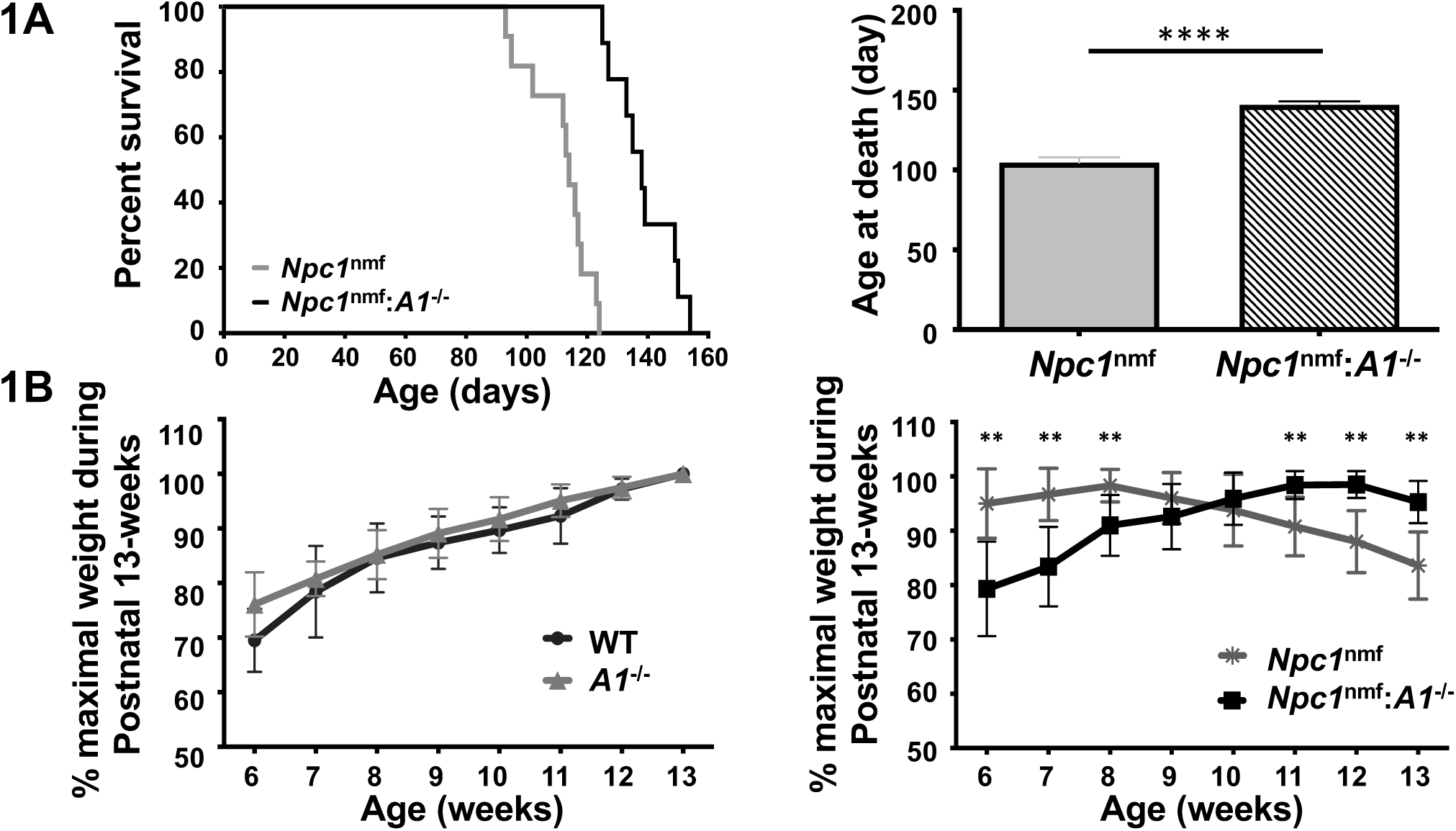
Effect of *A1*^-/-^ on the life span and weight loss of *Npc1^nmf^* mice. **A. *A1*^-/-^ increases life span of *Npc1*^nmf^ mice by 34%**. Median survival (left panel) for *Npc1nmf* mice and for *Npc1^nmf^:A1*^-/-^ mice is 113 days and 138 days respectively. Mean survival (right panel) is 102 days and 137 days, respectively. N=18 mice for *Npc1^nmf^* and N=16 for *Npc1*^nmf^:*A1*^-/-^ mice. Equal numbers of male and female mice were used, and the procedure described in (46) was adopted to define death of the *Npc1^nmf^* mouse. The *p*-value for survival curves is <0.0001. **B. *A1*^-/-^ delays weight loss of *Npc1^nmf^* mice (right panel) but not WT mice (left panel)**. Weight measurement began at 6 weeks of age. Data are expressed as percent maximum weight during the first 13 weeks. N=10 mice per group with equal numbers of males and females evaluated. Error bars indicate 1 SEM. In the right panel, except for weeks 9 (*p* = 0.2) and 10 (*p* = 0.4), the *p*-value for each week is <0.003. In the left panels, there are no significant differences.

Overall, *A1*^-/-^ increased *Npc1^nmf^* mutant mouse lifespan by 34%. In control experiments, no spontaneous deaths occurred in either WT mice or *A1*^-/-^ mice. We next evaluated the effect of *A1*-deficiency on the weights of *Npc1^nmf^* mice, starting at 6 weeks of age. Results (**Fig. 1B; right hand panel**) show that *Npc1^nmf^* mice start to lose weight at around 9 weeks of age. In contrast, mice lacking *A1* gene did not begin to lose weight until 13 weeks of age. Thus, the lack of *A1* delayed the onset of weight loss observed in *Npc1^nmf^* mice. In control experiments (**Fig. 1B; left hand panel**) at 6 weeks of age, *A1*^-/-^ mice weighed slightly less than WT mice, but at 8 weeks of age or older, the difference in weights between these 2 genotypes disappeared.

### *A1*^-/-^ reduced foam cell pathology in liver/spleen and Purkinje neuron loss in *Npc1^nmf^* mice

Niemann-Pick disease exhibits accumulation of large foamy macrophages in various tissues. To determine if *Acat1* inhibition had any effect on the foam cell pathology in *Npc1^nmf^* mouse, we isolated tissues from WT, *A1^-/-^, Npc1^nmf^*, and *Npc1^nmf^:A1^-/-^* mice at 80 days (P80) of age, and performed histological staining analyses. When compared to *Npc1*^*nm*f^ mice, *Npc1^nmf^:A1^-/-^* mice have significantly reduced foam cell pathology in liver (**Fig. 2A; right panels; top vs. bottom**) and spleen (**Fig. 2B; right panels; top vs. bottom**). In lung, however, foam cell pathology in these two genotypes are comparable (**Fig. 2C; right panels; top vs. bottom**). The result of the control experiments confirmed that neither WT mice nor global *A1*^-/-^ mice exhibit significant foam cell pathology in any of these three tissues (**Fig. 2A-C; left panel; top vs. bottom**). Previous studies in mutant NPC animals have demonstrated that extensive

**Fig. 2.**
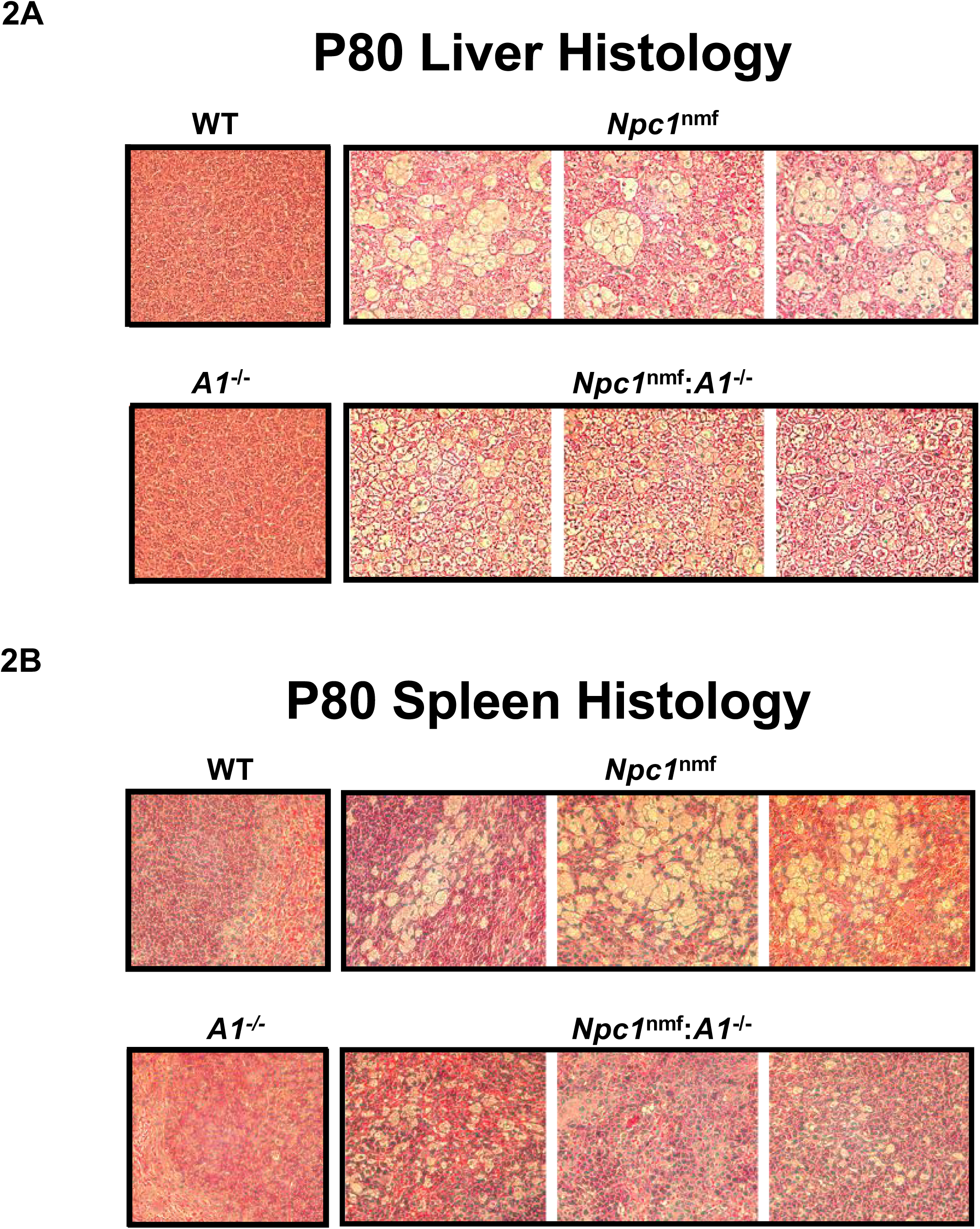
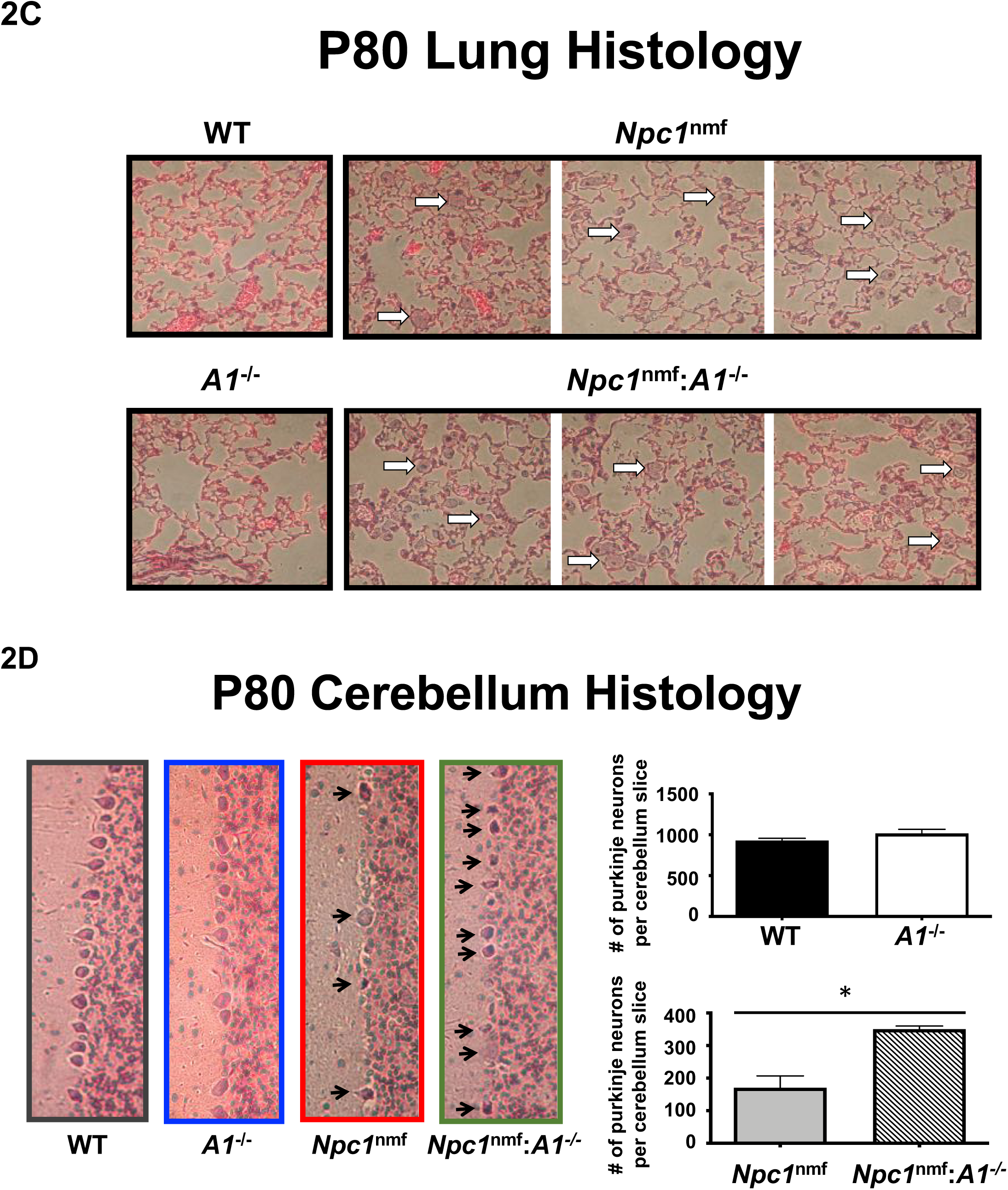
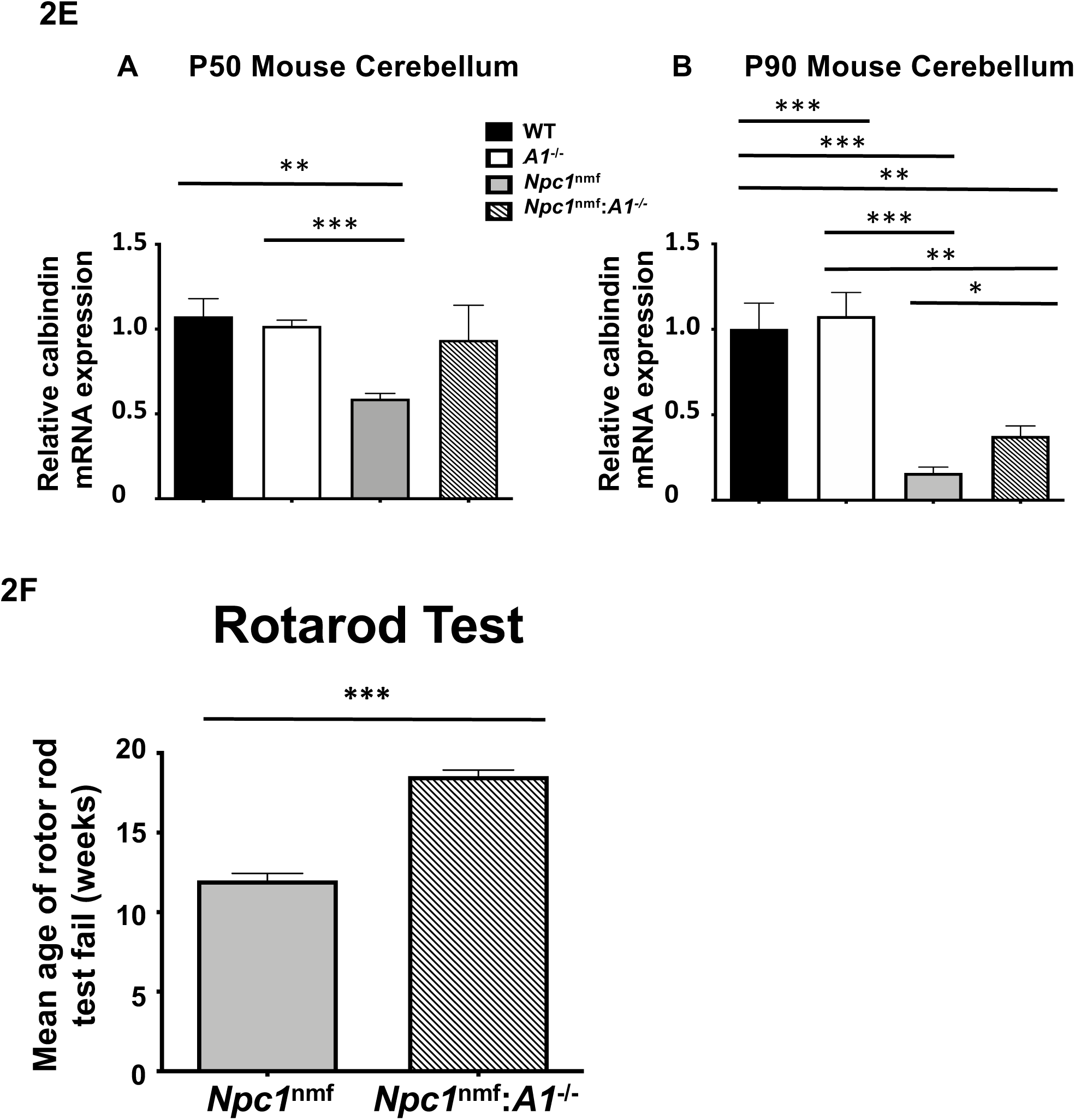
Effect of *A1*^-/-^ on cellular pathology, neuronal loss, and motor deficits in *Npc1*^nmf^ mice. **A-C**. *A1*^-/-^ improves macrophage foam cell pathology in liver (**A**), in spleen (**B**), but not in lung (**C**) of the *Npc1*^nmf^ mice. Tissues shown were isolated from mice at P80, fixed, sectioned, and processed for H& E staining. Lungs were perfused through the trachea. All mages were collected at the same magnification (40x). Results are representative of the 3 mice per group. In 2C, arrows point at foam cells. **D**. *A1*^-/-^ reduced Purkinje neuron death in *Npc1*^nmf^ mice, without affecting Purkinje neuron numbers in WT mice. Representative images of the cerebellum in the WT, *A1*^-/-^, *Npc1*^nmf^, and *Npc1^nmf^*: *A1*^-/-^ mice were collected at the same magnification (10x) and highlight the relative number of Purkinje neurons in each case. The relative number of Purkinje neurons (indicated by arrows in the images) was quantitated by counting in two separate cerebellar lobes. N=2 to 4 animals per group. Error bars indicate 1 SEM. For the right bottom panel, *p* <0.05. **E**. *A1*^-/-^ reduces the loss of calbindin expression in *Npc1*^nmf^ mouse cerebellum without affecting calbindin expression in WT mouse cerebellum. Cerebellar tissue was isolated from the brains of P50 (left panel) and P90 mice (right panel) and calbindin mRNA levels were measured by RT-PCR, with GAPDH also measured for normalization. Tissues were collected from 3 mice of each genotype. **F**. *A1*^-/-^ prevents the decline in motor skill in the *Npc1*^nmf^ mouse. Mice were tested each week in 3 consecutive trials on a rod rotating at a constant speed (24 rpm) for up to 90 s per trial. The fail time was defined as the age at which the mouse failed to stay on the rod at least 10 s during one of the 3 trials or froze on the rotarod and did not move. N=13 mice for *Npc1*^nmf^ and N=11 mice for *Npc1^nmf^*: *A1*^-/-^. Comparable numbers of male and female mice were evaluated. The *p-*value is <0.001.

Purkinje neuron cell death occurs prior to the death of the animals. To evaluate the effect of *A1*^-/-^ on *Npc1^nmf^* Purkinje neurons, cerebellum was isolated from P80 mice brain, and the number of Purkinje neurons were counted after histochemical staining of thin slices. As shown in **Fig. 2D**, when compared to values that found in WT mice and in *A1*^-/-^ mice, *Npc1^nmf^* mice had less than 20% residual Purkinje cells remaining, while in the *Npc1^nmf^:A1^-/-^* mice, the numbers of residual Purkinje neurons remaining was more than 30% of WT values (**Fig. 2D; right panels**). To validate the results shown in Fig. 2D, we isolated cerebellar tissue from the brains of P50 and P90 mice, and compared the levels of calbindin mRNA, as calbindin expression is specific to Purkinje neurons in the cerebellum. The results of P50 cerebellum (**Fig. 2E; A**), calbindin mRNA levels in WT and *A1*^-/-^ mice were comparable (first 2 bars on left). In *Npc1^nmf^* mice (the 3^rd^ bar from left), the expression was significantly reduced by 55% of the values found in WT and *A1*^-/-^ mice, but the expression was almost completely restored in *Npc1^nmf^:A1^-/-^* mice (4^th^ bar from left) to about 90% of values found in WT and *A1*^-/-^ mice. The results obtained from the P90 cerebellum (**Fig. 2E; B**) show that calbindin expression in *Npc1^nmf^* mice was drastically reduced to about 10% of values found in WT and *A1*^-/-^ mice, but was partially restored in *Npc1^nmf^:A1^-/-^* mice, to about 30% of values found in WT and *A1*^-/-^ mice. Overall, these results demonstrate that in the *Npc1^nmf^* mouse model, *A1* gene ablation significantly decreases the loss of Purkinje neurons in brain.

### *A1*^-/-^ ameliorated the motor deficits and behavior of *Npc1*^nmf^ mice

Mutant *Npc1^nmf^* mice exhibit a decline in their motor performance, beginning at 11 weeks of age (46). We monitored the motor performance of WT, *A1^-/-^, Npc1^nmf^*, and *Npc1^nmf^*:*A1^-/-^* mice beginning at 9 weeks of age by using a rotarod test. The results show that for *Npc1^nmf^* mice, the mean age of failing off or not running on the rotarod test occurred at around 12 weeks of age, whereas for *Npc1^nmf^:A1^-/-^* mice, the mean age of failing occurred at 18 weeks of age (**Fig. 2F**). Results from control experiments show that at the same age, neither WT mice nor *A1*^-/-^ mice fail the rotarod test. Since the rotarod test does not assess muscle strength per se, these results do demonstrate that *A1* gene ablation improves sensorimotor coordination and behavior in the *Npc1^nmf^* mice.

### Cholesterol content, cholesterol esterification, and cholesterol distribution in mouse embryonic fibroblasts (MEF) from WT, *A1^-/-^, Npc1^nmf^*, and *Npc1^nmf^:A1^-/-^* mice

To determine whether *A1^-/-^* alters the total free cholesterol content in *Npc1*^*nm*f^ mouse brain, we first isolated whole brains from WT, *A1^-/-^, Npc1^nmf^*, and *Npc1^nmf^*:*A1^-/-^* mice at postnatal day 90 (P90) and measured their total free (unesterified) cholesterol content. The results show that the cholesterol content in these samples are comparable to one another (**Fig. 3A**).

**Fig. 3.**
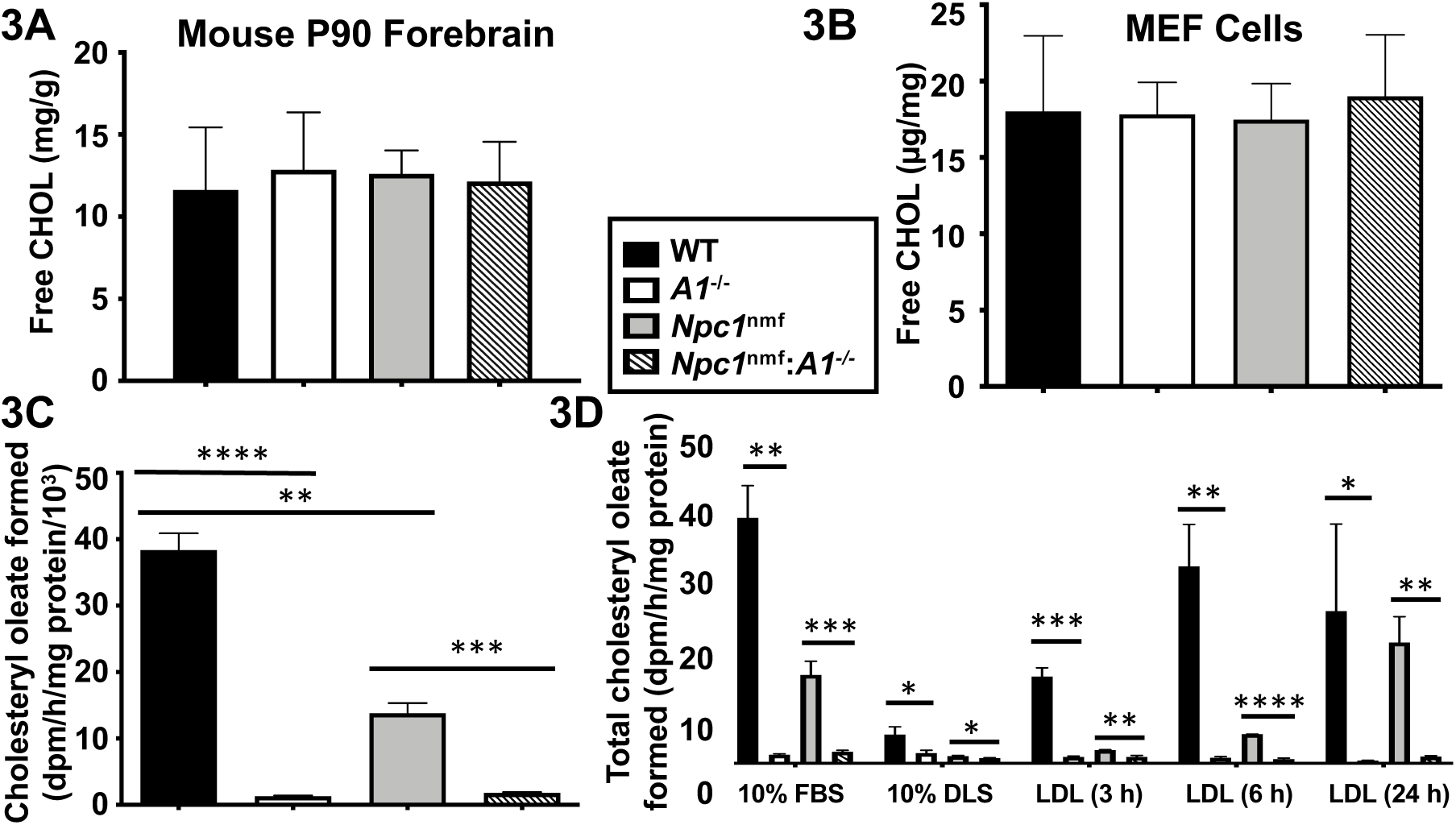
Cholesterol content and cholesteryl ester biosynthesis. **A. Free cholesterol content in P90 WT, *A1*^-/-^, *Npc1*^nmf^, and *Npc1*^nmf^ :*A1*^-/-^ mouse forebrain**. Tissues were homogenized in chloroform: methanol 2:1, filtered by Whatman filter paper, dried under nitrogen then resuspended in methanol. Free cholesterol content was determined in triplicate by using Wako’s Free Cholesterol E kit. N= 5 mice (3 male and 2 female) per group. Error bars indicate 1 SEM. **B. Free cholesterol content in mouse embryonic fibroblasts (MEFs)**. MEF were seeded in 6-well dishes at 200,000 cells/well and grown in DMEM plus 10% serum to near confluence. After three washes with PBS, cells were scraped off the dish to form suspensions in PBS and were used for protein measurement and lipid extraction by chloroform/methanol. Cholesterol content was determined as described in part A above. Data shown is from 3 platings per genotype. **C. Cholesteryl ester biosynthesis in MEFs continuously grown in lipoprotein containing medium**. Cells were grown as described in part B above. Cholesteryl ester biosynthesis in intact cells was described in (103). Data shown is from 3 platings per genotype. **D. Cholesterol ester biosynthesis in MEFs grown in cholesterol containing medium (10% FBS), cholesterol free medium (10% DLS), or 10% DLS in response to LDL feeding for 3 h, 6 h, or 24 h**. Human LDL and delipidated fetal bovine serum were prepared as described (103).

In the central nervous system, however, the bulk of cholesterol exists in the membranes of a variety of cell types, including the glial-derived myelin sheath, making measurements of bulk cholesterol less informative when considering the effects of NPC1 deficiency, as well as the effects of ACAT1 inhibition on membrane cholesterol distribution at the cellular level. Parallel cultures of primary fibroblasts isolated from normal mice and from mice with single mutations have been used extensively as a model system to investigate the effects of single-gene mutations at the biochemical level. We therefore used mouse embryonic fibroblasts (MEFs) to address this question. We isolated MEFs from WT, *A1^-/-^, Npc1^nmf^*, and *Npc1^nmf^:A1^-/-^* mice, grew them in DMEM with 10% serum in monolayers until confluent, and then harvested the cells for the analysis of total free cholesterol content. Results (**Fig. 3B**) show that the free cholesterol content in these four cell types are very similar, demonstrating that under the condition employed, neither NPC1 mutation nor ACAT1 blockage significantly affects the total cellular free cholesterol content.

We next measured relative cholesterol ester biosynthesis rates in these cells by feeding labelled 3H-oleate to the intact cells for 20-min and then measuring the ^3^H-cholesteryl oleate that was produced. The results (**Fig. 3C**) show that when grown in lipoprotein-containing medium (10% serum), WT cells exhibit an ample cholesterol ester biosynthesis rate. As expected, in cells without ACAT1 (i.e., *A1^-/-^* cells and *Npc1^nmf^:A1^-/-^* cells), the cholesterol ester biosynthesis rate was only 2% that of WT cells. The residual amount of cholesteryl oleate in ACAT-deficient cells formed are perhaps due to the presence of ACAT2 (48). In *NpC1^nmf^* cells, the rate was around 34% that of WT cells (**Fig. 3C**). When these four cell types were grown in 10% delipidated serum medium (i.e., medium devoid of exogenous cholesterol), the cholesteryl ester biosynthesis rate in WT cells decreased drastically; these decreases also occurred in the other three cell types (**Fig. 3D; lanes 5-8**); whereas, in contrast, when low-density lipoproteins (LDL) were added to the delipidated serum (DLS) medium for 3 h, 6 h, or for 24 h (indicated as LDL (3 h), LDL (6 h) or LDL (24 h) below the X-axis in **Fig. 3D**), cholesteryl ester biosynthesis rate was restored in WT cells in a time-dependent manner. Adding LDL also increased cholesteryl ester biosynthesis rate in mutant *Npc1^nmf^* cells, but this increase was not nearly as what was observed in WT cells (**Fig. 3D; comparing 3^rd^ bars vs. 1^st^ bars**). These results show that within the 3-6 h period, mutant *Npc1^nmf^* cells fail to efficiently utilize LDL-derived cholesterol for esterification, indicating the importance of NPC1 in delivering cholesterol from endosomes to the ER. As expected, in *A1^-/-^* cells or in *Npc1^nmf^:A1^-/-^* cells, within the 3-6 h time frame, the LDL-dependent increase in cholesterol ester biosynthesis rate was abolished (**Fig. 3D; comparing 2^nd^ bars and 4^th^ bars vs. the 1^st^ bars**). When cells were exposed to LDL for 24 h, a large increase in cholesterol ester biosynthesis rate occurred in *Npc1^nmf^* cells but not in cells without *A1* (**Fig. 3D**; comparing last 4 bars at the LDL (24 h) time point). Our earlier results (**Fig. 3C**) show that, when maintained in the lipoprotein-containing medium at a steady state, *Npc1^nmf^* cells exhibit a residual cholesterol ester biosynthesis rate of about 34% compared to WT cells. These results support the notion (described in the Introduction) that, in addition to using cholesterol derived from the NPC-containing late endosomes for esterification, cells can also use other cholesterol sources for esterification at the ER, in an NPC-independent manner.

ACAT1 plays a key role in cholesterol storage via its conversion of cholesterol to cholesterol esters. In both WT and *Npc1^nmf^* MEFs, preventing cholesterol storage by inhibiting ACAT1 may cause significant alteration(s) in cholesterol distribution among various membrane organelles. To test this possibility, we performed intact cell staining with filipin, a naturally fluorescent small molecule that binds to cholesterol, and then viewed the labeled cells with spinning disc confocal fluorescence microscopy. The results show that in WT and *A1^-/-^* cells (**Fig. 4A; 1^st^ and 2^nd^ columns**), the cholesterol-rich domains, as represented by areas with strong filipin staining (**1^st^ row**), were mostly located in peripheral regions of the cytosol, whereas in mutant *Npc1^nmf^* cells (**Fig. 4A; 3^rd^ column**), the cholesterol-rich domains were primarily peri-nuclear in their localization. It was previously shown that LE/LYS exhibit non-random, bidirectional movements between the nucleus region and the cell surface; when mutant NPC1 cells were laden with cholesterol and other lipids, the movements of LE/LYS towards the cell periphery became sluggish, presenting cholesterol-rich particles primarily to the perinuclear location (49). The results shown in Fig. 4A; 3^rd^ row confirms these findings. Importantly, in mutant *Npc1^nmf^* cells, *A1^-/-^* caused most of the cholesterol-rich particles to become primarily peripherally located (**Fig. 4A; the 4^th^ column**). We also calculated total cell fluorescence intensity of filipin per cell and found that the values among these four cell types were comparable (**Fig. 4B**). Overall, these results imply that *A1^-/-^* corrected the late endo/lysosomal cholesterol sequestration defects seen in the mutant NPC1 cells without significantly altering the total cholesterol content in these cells. We also noted that when compared to WT cells, *A1*^-/-^ cells contained more cholesterol-rich domains that are scattered throughout the cytoplasm (**Fig. 4A; the 2^nd^ column**), suggesting that in WT cells, *A1*^-/-^ causes significant cholesterol accumulation in certain internal membrane organelle(s). The nature of these cholesterol-rich internal organelles is unknown at present.

**Fig. 4.**
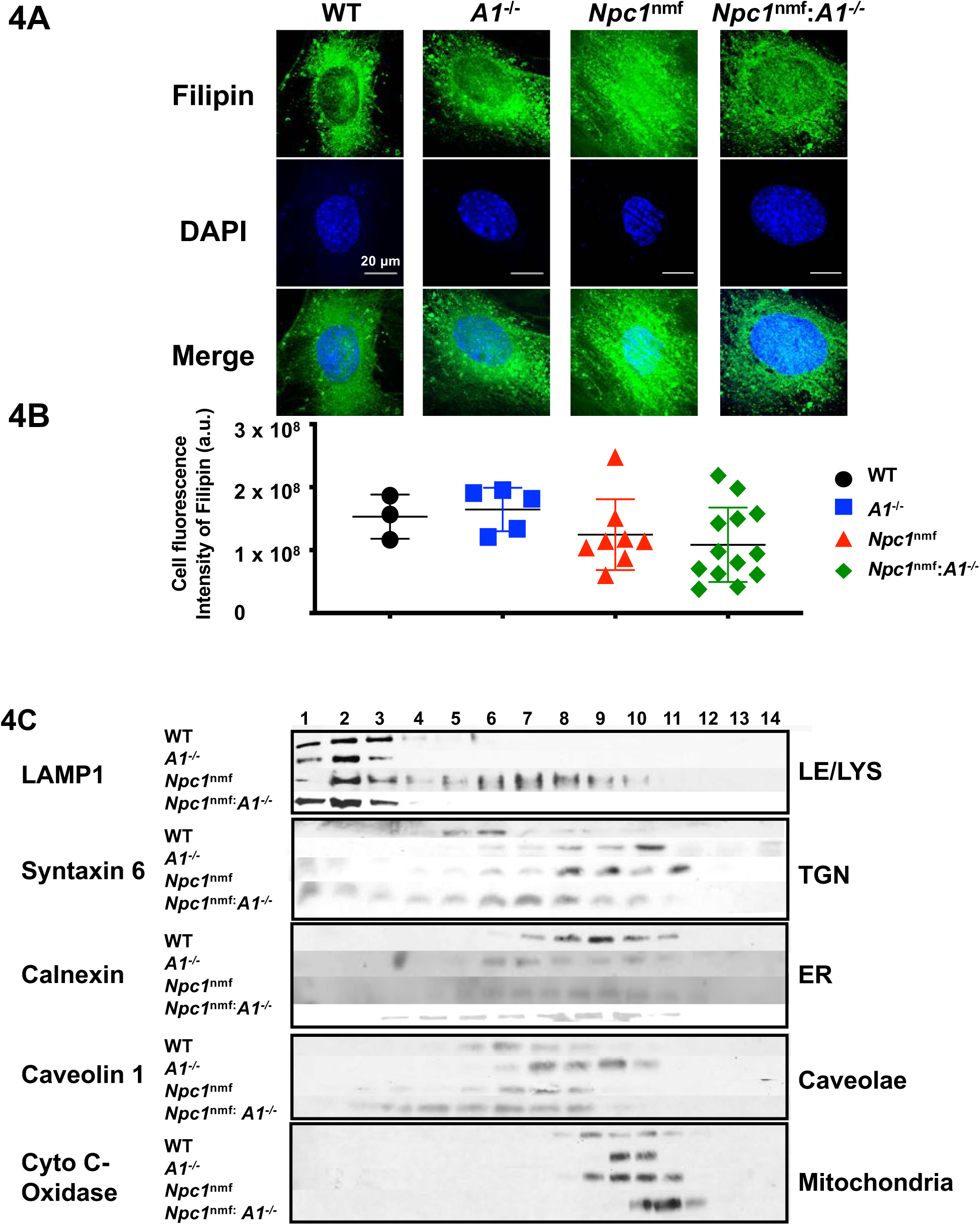
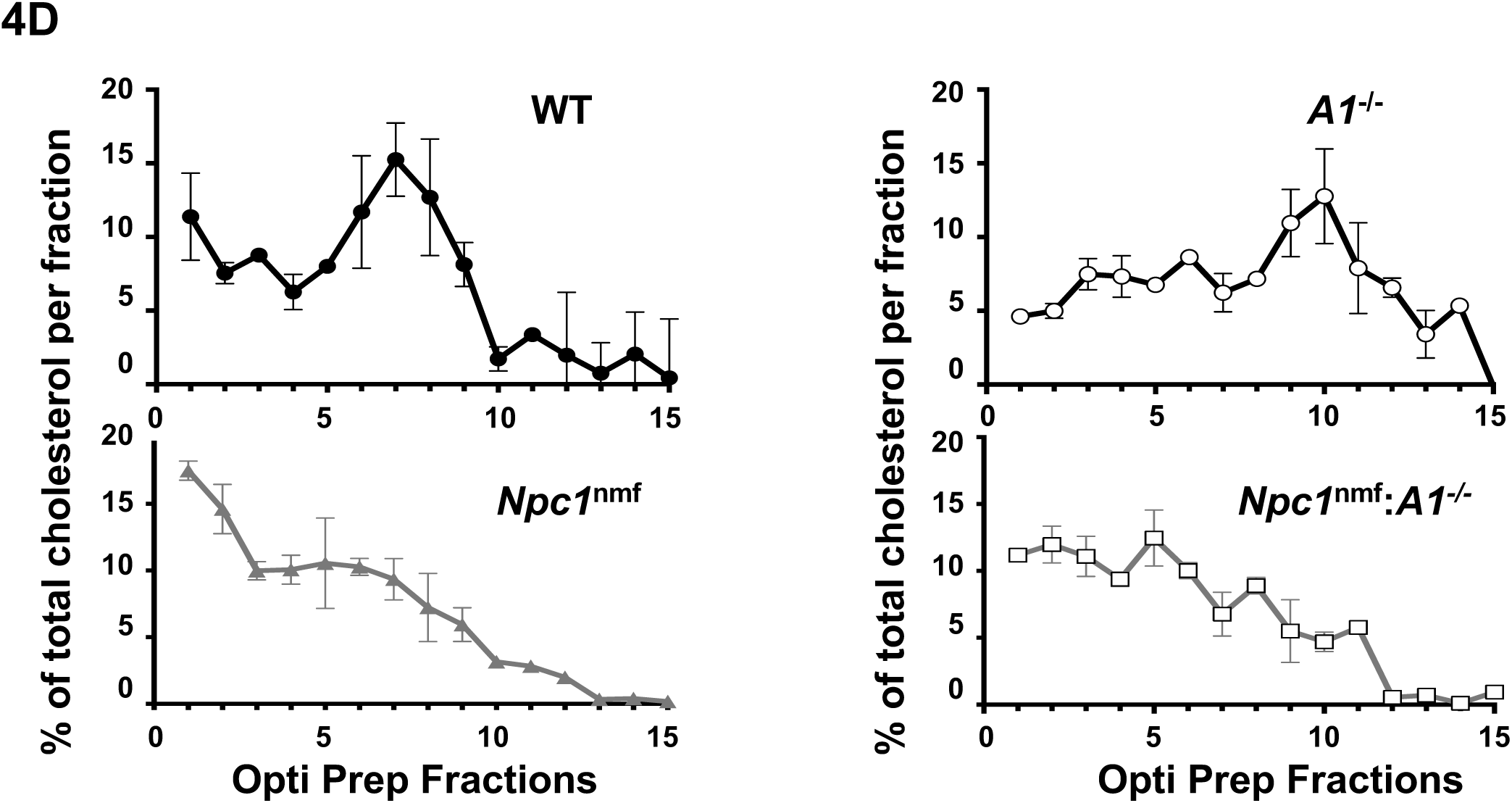
Cholesterol distribution in mouse embryonic fibroblasts (MEFs). **A**. MEFs from WT, *A1*^-/-^, *Npc1*^nmf^, and *Npc1*^nmf^ :*A1*^-/-^ mice were seeded on poly-d-lysine treated glass coverslips in 12-well plates. Filipin staining was carried out according to (104). **B**. Total fluorescence intensity per cell. Fluorescence was quantified using NIH Image J. Each point represents the value obtained for an individual cell. **C**. Localization of subcellular organelles after cell fractionation. Antibodies for specific protein markers (listed on far left) were used to localize specific organelles (listed on right). **D**. Cholesterol distribution in various subcellular organelles. For each cell type, cells from a single 15-cm dish were grown in medium plus 10% serum until confluent. Cell homogenization and OptiPrep density gradient ultracentrifugation, were carried out as described (22). Fourteen fractions were collected from each sample, with each fraction analyzed in triplicate for cholesterol content using a Wako kit. The values reported are normalized to the total cholesterol content present in all fractions. Data are means +/-1 SD from 2 separate experiments.

To examine the membrane cholesterol distribution in these four cell types with another method, we took a biochemical approach, and performed subcellular fractionation of post-nuclear cell homogenates prepared from the MEFs of these four different genotypes, using OptiPrep density gradient ultracentrifugation. This method produces partial separation of various membrane organelles, based primarily on their buoyant densities (50), (51), (22). After OptiPrep fractionation we analyzed the distribution of various membrane organelles in the fractions by performing Western blot analyses of protein markers for LE/LYS, TGN, ER, caveolae, and mitochondria (**Fig. 4C**). We also analyzed cholesterol content in each fraction (**Fig. 4D**).

Western analyses (**Fig. 4C**) show that in all four cell types, LE/LYS (LAMP1 positive) were tightly enriched in fractions #1 to #3 (with light density) and the mitochondria (cytochrome C oxidase positive) were tightly enriched in fractions #9 to #11 (with heavy density). Mutant *Npc1^nmf^* cells, but not *Npc1^nmf^:A1^-/-^* cells, exhibited additional LAMP1 positive signals in fractions #5 to #9. The ER (calnexin positive) exhibited a broader range of densities but was similarly enriched in #6 to #10 in all cell types. The PM (caveolin 1 positive) was enriched in fractions #5 to #8 in WT cells (with medium density), but in *A1^-/-^* cells was enriched in later fractions (#7 to #9). This result suggests that *A1*^-/-^ may cause an increase in buoyant density of the PM. Similar to the WT cells, in mutant *Npc1^nmf^* cells and in *Npc1^nmf^*:*A1*^-/-^ cells, the PM were enriched in fractions #5 to #8. However, the PM in mutant *Npc1^nmf^:A1*^-/-^ cells exhibited a broader range in buoyant densities. These results show that except for the PM in *A1*^-/-^ cells and the abnormal LAMP1 positive signal in mutant *Npc1^nmf^* cells, the buoyant densities of LE/LYS, mitochondria, ER, and PMs in these four cell types are comparable. In contrast, the syntaxin 6 rich-fractions (**Fig. 4C; 2^nd^ row**) exhibited large variation in densities: in WT cells, they are enriched in fractions #5 to #6; in *A1*^-/-^ cells, in fractions #8 to #10; in *Npc1^nmf^* cells, in fractions #8 to #11, in *Npc1^nmf^:A1^-/-^ cells*, in fractions #5 to #9. The results of cholesterol content analyses in the OptiPrep fractions (**Fig. 4D**) show that as expected, in WT cells, cholesterol was highly enriched in the PM fraction (#5 to #8), and membranes with slightly heavier densities, including the ER membranes (#7 to #9), whereas the mitochondrial membranes had much less cholesterol content. In *A1*^-/-^ cells, membranes with heavier densities, including the ER (#7 to #9) and mitochondria (#9 to #11), may be enriched in cholesterol. However, this interpretation is not definitive, because in the *A1^-/-^* cells, the PM fractions increased in buoyant densities, such that they overlapped significantly with the ER fractions (#7 to #9) (**Fig. 4C; 4^th^ row**). In mutant *Npc1^nmf^* cells, as expected, cholesterol was highly enriched in the light density late endo/lysosomes (LE/LYS) fraction (fractions #1 to #3). In the mutant *Npc1^nmf^:A1^-/-^* cells, the Golgi-like membranes (#6 to #9), the PM-like membranes (#5 to #8), and the ER-like membranes (#7 to #9), all become relatively enriched in cholesterol. These results corroborate with the cholesterol distribution in intact cells (shown in Fig. 4A), and support the interpretation that while in *Npc1^nmf^* cells cholesterol is mostly sequestered within LE/LYS, with the deletion of the *A1* gene in *Npc1^nmf^:A1^-/-^ cells* most of the sequestered cholesterols appears redistributed to various other membrane organelles, including Golgi, ER, PM, and mitochondria.

### Effects of *A1^-/-^* on syntaxin 6 and golgin 97 localization in intact mutant *Npc1^nmf^* MEFs

The result presented in **Fig. 4C** (2^nd^ row) showed that the syntaxin 6-rich membranes isolated from the four different MEFs exhibited large variation in buoyant densities. Syntaxin 6 binds to cholesterol (52), and is one of the t-SNARE proteins present in vesicles that participate in various membrane fusion events. The syntaxin 6-rich vesicles move dynamically between various membrane organelles. In normal cells, most of the syntaxin 6 signal is found at the TGN (53). The TGN is rich in cholesterol content (54), and plays key roles in transporting proteins and lipids to various other membrane compartments, including the PMs and endosomes. In mutant NPC cells, the TGN fails to receive LDL-derived cholesterol from the late endosomes (22). This deficiency causes syntaxin 6 to be mislocalized from the TGN, and instead it exhibits an abnormal, scattered cytoplasmic pattern (25). Importantly, treating mutant NPC1 cells with cholesterol/cyclodextrin complex, or with high concentration of LDL for 24 h restores the syntaxin 6 signal to the typically one-sided, perinuclear Golgi localization pattern observed in normal cells (25). In mutant NPC cells, other SNARE proteins such as syntaxin 16, VAMP3, VAMP4, do not show cholesterol-sensitive localization patterns (25). Based on these previous findings, as well as results presented in **Fig. 4B**,**C**,**D**, we postulated that *A1^-/-^* may correct the mis-localization pattern of syntaxin 6 observed in mutant *Npc1^nmf^* MEFs. To test this possibility, we performed immunofluorescence confocal microscopy in fixed, intact cells using antibodies specific for syntaxin 6. The results (**Fig. 5A**) show that, in WT cells and in *A1^-/-^* cells most of the syntaxin 6 signal was highly polarized to only one side of the space adjacent to the nucleus. This pattern is a typical Golgi distribution pattern found in mammalian cells (55). In contrast, in mutant *Npc1^nmf^* cells (**Fig 5A**), a significant portion of the syntaxin 6 signal was distributed in scattered cytoplasmic vesicular structures; these structures were located at the space around both sides of the nucleus. This result confirms the findings by Reverter *et al*. (25). Importantly, A1 deletion (*A1^-/-^*) in mutant NPC cells (**Fig 5A**) largely restored the syntaxin 6 distribution pattern back to the one-sided, peri-nuclear pattern observed in WT and *A1^-/-^* cells. To quantitate the difference of syntaxin 6 positive signals observed in the MEFs of these four different genotypes, we adopted the procedure developed by Mitchel *et al*. (55), by measuring the “Reflex angle”, defined as the angle subtended by the edges of the syntaxin 6-positive signals in the confocal images, using the center of the nucleus (DAPI positive signal) as the vertex. The results (**Fig. 5B**) show that for WT and *A1^-/-^* cells, the reflex angle was very similar : 119^°^ or 126^°^ respectively; for mutant *Npc1^nmf^* cells, the reflex angle was much larger (346^°^). For mutant *Npc1^nmf^*:*A1^-/-^* cells, the reflex angle was largely restored, to 160^°^.

**Fig. 5.**
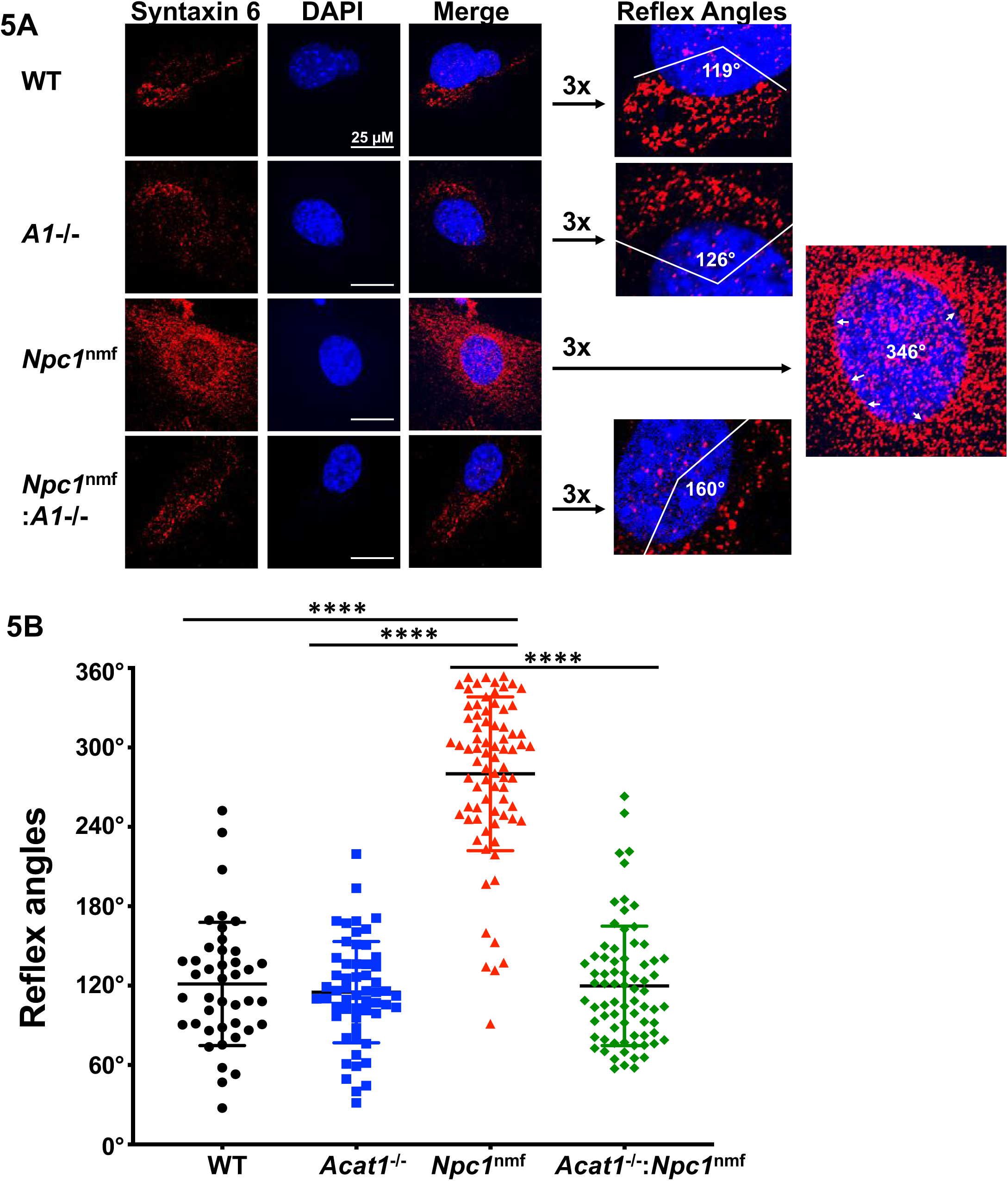
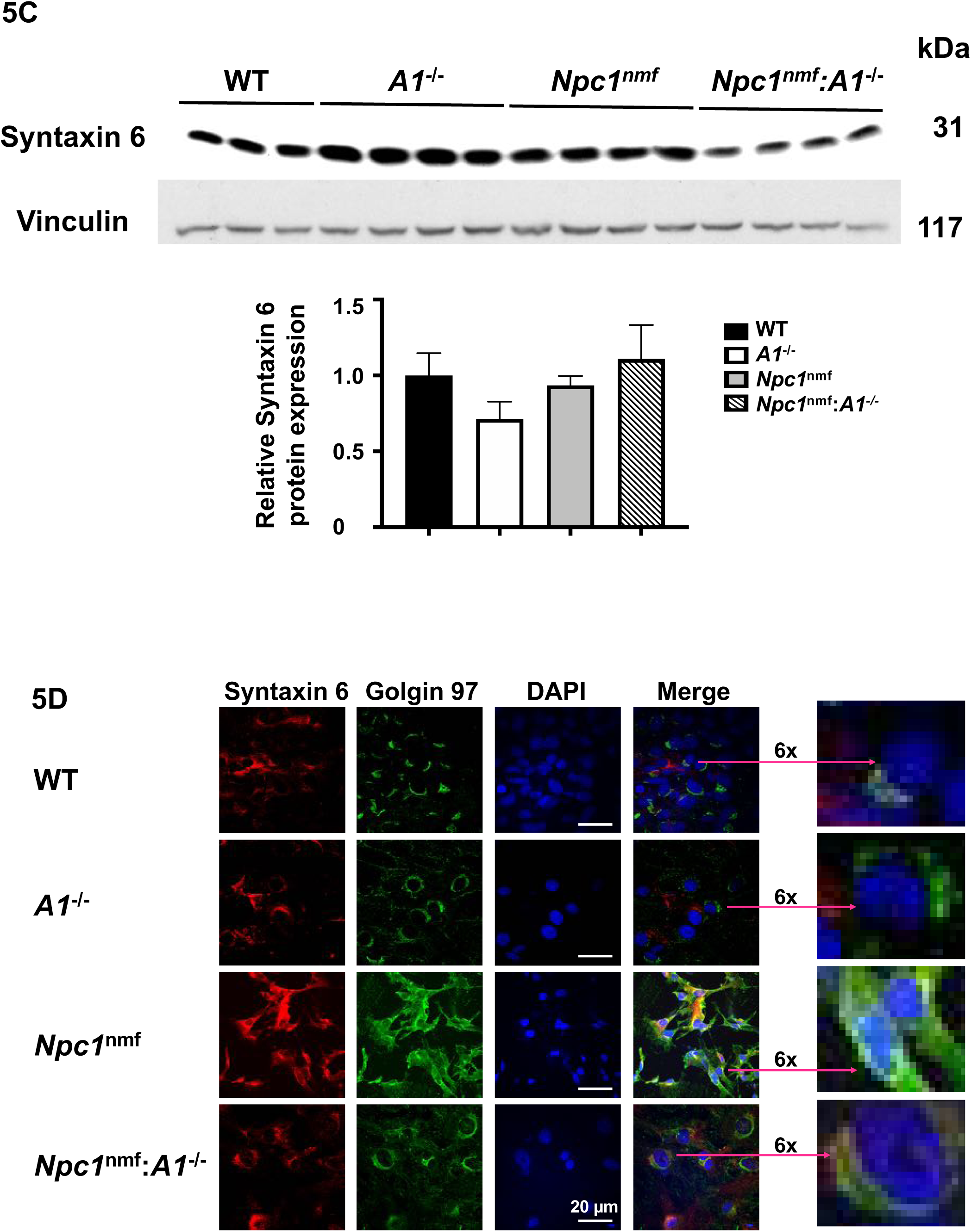
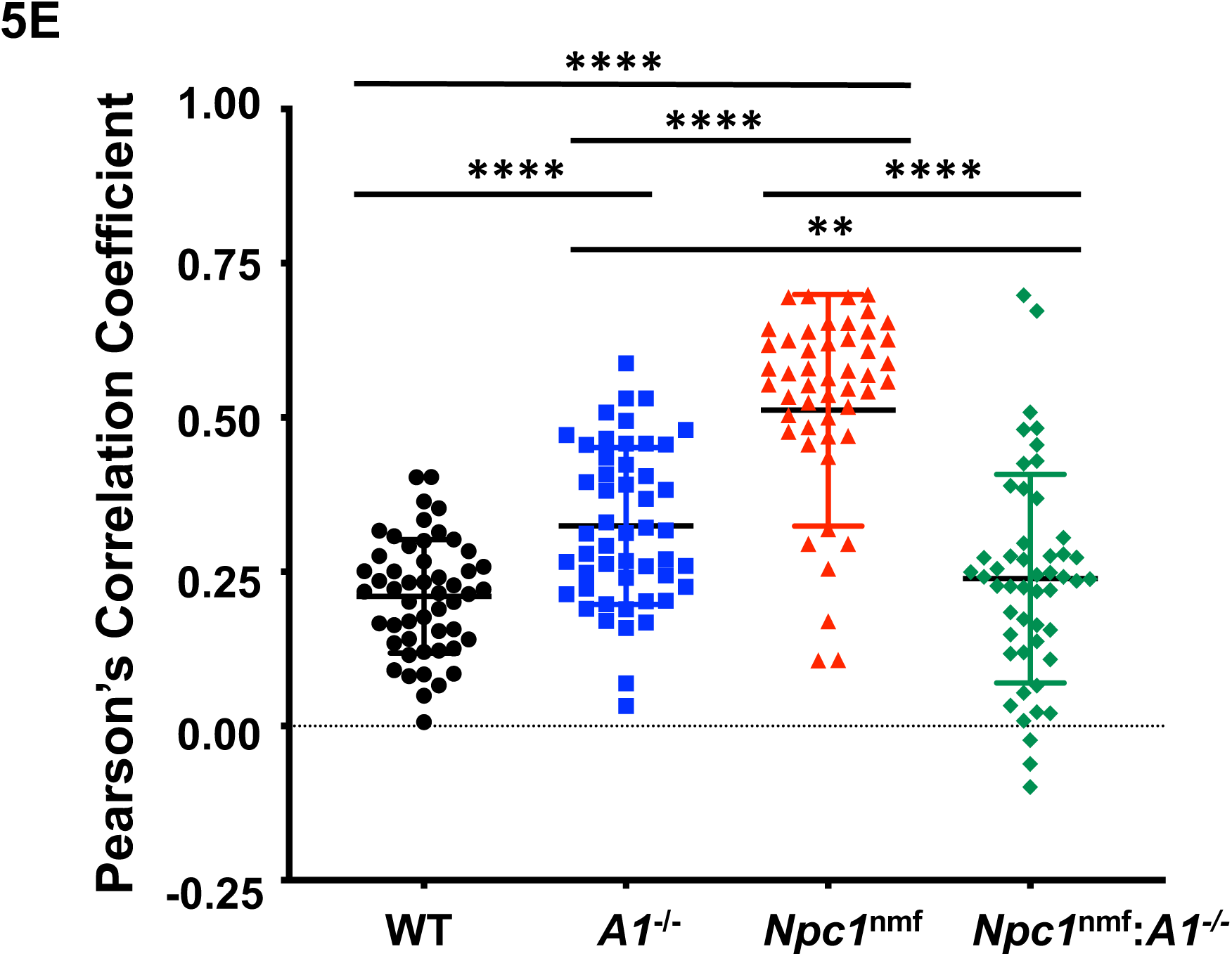
Localization of syntaxin 6 and golgin 97 in intact mouse embryonic fibroblasts (MEFs). **A**. The methods used for the detection of fluorescence-labeled **syntaxin 6 in** MEFs are described in the Methods. Fiji-ImageJ software was used to calculate the “Reflex angle”, which is indicated in the images on the right. **B**. The Reflex Angle of individual cells from WT, *A1*^-/-^, *Npc1*^nmf^, and *Npc1*^nmf^ :*A1*^-/-^ mice. Between 50 to 70 cells were measured for each genotype. **C. Relative syntaxin 6 protein content in lysates of MEFs**. Cells were seeded in 60 mm culture dishes and grown in DMEM plus 10% serum until confluent. Then 200 µl of 10% SDS were added per dish, and 150 µg of the solubilized protein was loaded per lane for Western blot analyses. The vinculin signal was used as the loading control. Error bars indicate 1 SEM. **D. Double immunofluorescence of syntaxin 6 and golgin 97 in fixed, intact MEFs**. The conditions used were the same as described in part A above. **E. Degree of apparent colocalization between syntaxin 6 and golgin 97 in MEFs**. Nikon NIS-Element AR Imaging Software was used to create the Maximum Intensity Projection used to calculate the degree of colocalization, which is reported as Pearson’s correlation coefficient. Each data point represents the result from an individual cell, and images from more than 50 cells were collected for each genotype. GraphPad Prism’s One-way ANOVA multiple comparisons method was used for statistical analysis.

The results of control experiments (**Fig. 5C**) show that the level of syntaxin 6 protein in all four cell types examined were comparable. Together, these results show that *A1^-/-^* largely restores the mis-localization of syntaxin 6 observed in mutant *Npc1^nmf^* cells, without affecting the syntaxin 6 protein content.

To strengthen these results, we sought to examine the localization pattern of a second protein marker of the TGN. Golgins are long coiled-coil peripheral membrane proteins located mainly at the membrane surface of the TGN (56). In humans there are four golgins, each playing a distinct role in membrane protein transport events. Using specific antibodies against golgin 97 (GCC97), we performed double immunofluorescence experiments to compare the localization patterns of syntaxin 6 and golgin 97 in parallel cultures of four MEF cell types. The results show that, in WT cells (**Fig. 5D, 1^st^ row)**, golgin 97 exhibited a typical polarized Golgi distribution pattern. In *A1^-/-^* cells (**Fig. 5D, 2^nd^ row**), most of the golgin 97 exhibited a similar pattern to what was found in WT cells; with a small portion also appearing as part of small, punctate structures, perhaps as parts of the internal membrane organelles. In *Npc1^nmf^* cells **(Fig. 5D; 3^rd^ row**), golgin 97 became dispersed, and scattered within the space around the nucleus. This distribution pattern is clearly distinct from that observed in WT cells and in *A1*^-/-^ cells. Importantly, in *Npc1^nmf^*:*A1^-/-^* cells (**Fig. 5D; 4^th^ row**), the abnormal golgin 97 localization pattern in *Npc1^nmf^* cells was largely corrected. For clarity, enlarged (6-fold) versions of the merged images are provided (**Fig. 5D**). We next performed double immunofluorescence experiments and monitored the degree of apparent colocalization between syntaxin 6 and golgin 97 in these four cell types (**Fig. 5 D, E**). The results show that in WT and *A1^-/-^* cells, the apparent colocalization between syntaxin 6 and golgin 97 was relatively low (between 21% to 32%). Mutation in NPC1 caused the colocalization index to increase to 51%; while *A1 deletion (A1*^-/-^) in the mutant *Npc1^nmf^* cells reduced the % colocalization index back to 24%, a value comparable to those observed in WT and *A1^-/-^* cells. Together, these results shown in **Fig. 5A-E** suggest that mutation of NPC1 causes a certain portion of the TGN membrane to become deficient in cholesterol; this deficiency causes both syntaxin 6 and golgin 97 to become mislocalized from the rest of the TGN. Importantly, ACAT1 blockage in mutant NPC1 cells largely corrects the abnormal localization patterns of both syntaxin 6 and golgin 97.

### Effects of *A1^-/-^* on the levels of cation-dependent manose-6-phosphate receptor (CD-M6PR), and cathepsin D protein contents in mutant *Npc1^nmf^* MEFs

Syntaxin 6 is involved in the anterograde vesicular trafficking of M6PRs/lysosomal hydrolase complexes (57). The cation-dependent (CD) and cation-independent (CI) mannose-6-phosphate receptors (CD-M6PR and CI-M6PR) deliver newly synthesized lysosomal enzymes, which carry the mannose-6-phosphate signal, from the TGN to the late endosomes. The M6PRs then recycle back to the TGN for re-utilization, as reviewed in (58). Previously, Kobayashi *et al*. (59) showed that in mutant NPC cells, the CD-M6PR localization pattern was altered, from mainly residing at the TGN to mostlyly residing in the cholesterol-laden late endosomes. Ganley and Pfeffer (60) showed that in mutant NPC cells, cholesterol accumulation caused CD-M6PR to be missorted and rapidly degraded in late endo/lysosomes. Since we found that in mutant *Npc1^nmf^* cells, *A1^-/-^* rescued syntaxin 6 from mislocalization (**Fig. 5A**,**B**), we suspected that *A1^-/-^* may also affect the M6PR protein expression in mutant *Npc1^nmf^* cells. To test this possibility, we performed Western blot analyses to examine the levels of CD-M6PR protein in parallel cultures of MEFS of the four genotypes. The results (**Fig. 6A**) show that when compared to WT cells, *Npc1^nmf^* cells express CD-M6PR at levels less than 20% of values found in WT cells. In contrast, the *Npc1^nmf^:A1*^-/-^ cells expressed CD-M6PR at levels more than 2-fold that of WT cells. This result suggests that deleting *A1* was indeed more than sufficient to correct the low level of expression of the CD-M6PR protein observed in *Npc1^nmf^* cells. Interestingly, we also found that *A1^-/-^* cells expressed CD-M6PR level at 50% of values found in WT cells. At present, the significance of this finding is unclear at present.

**Fig 6.**
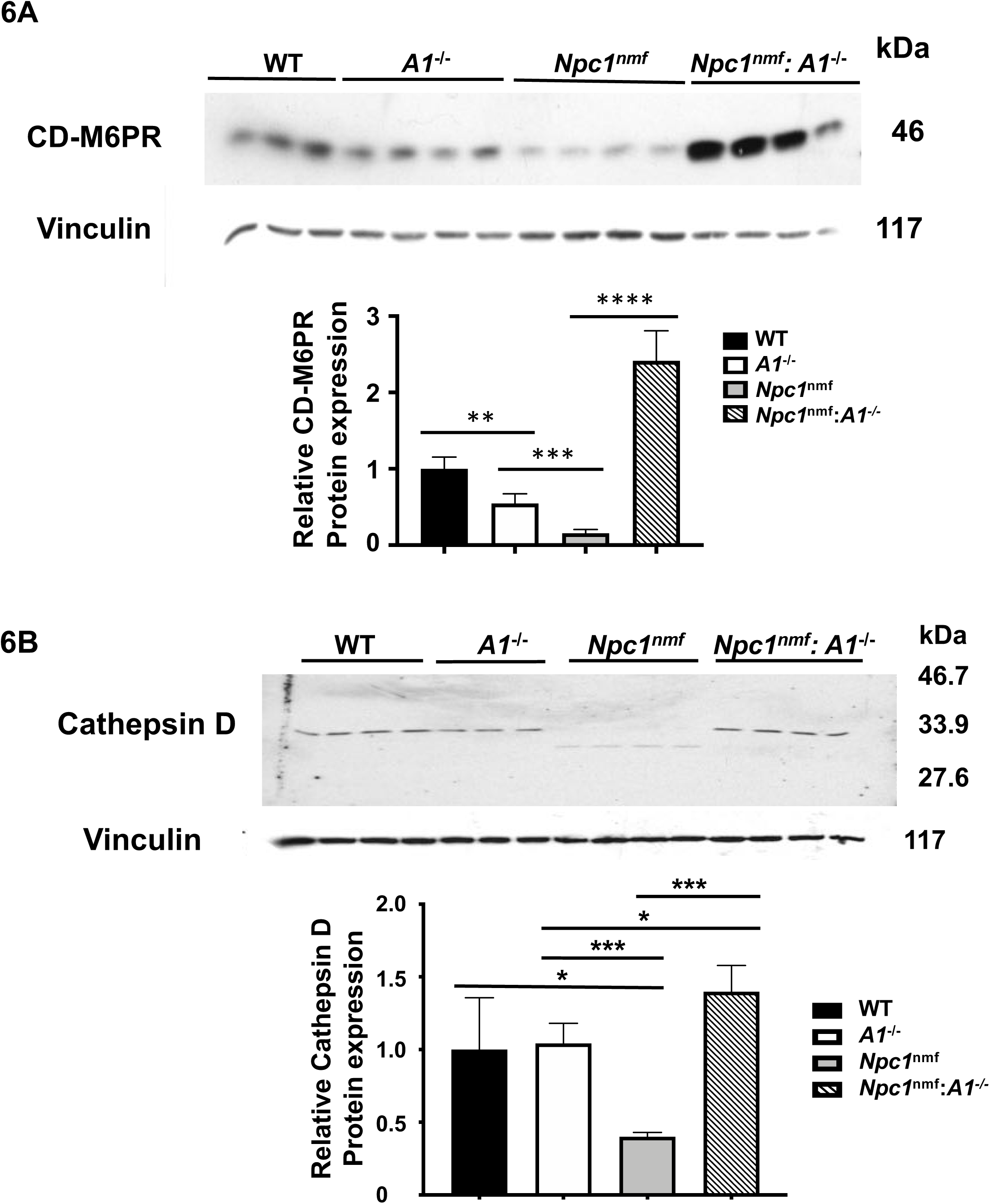
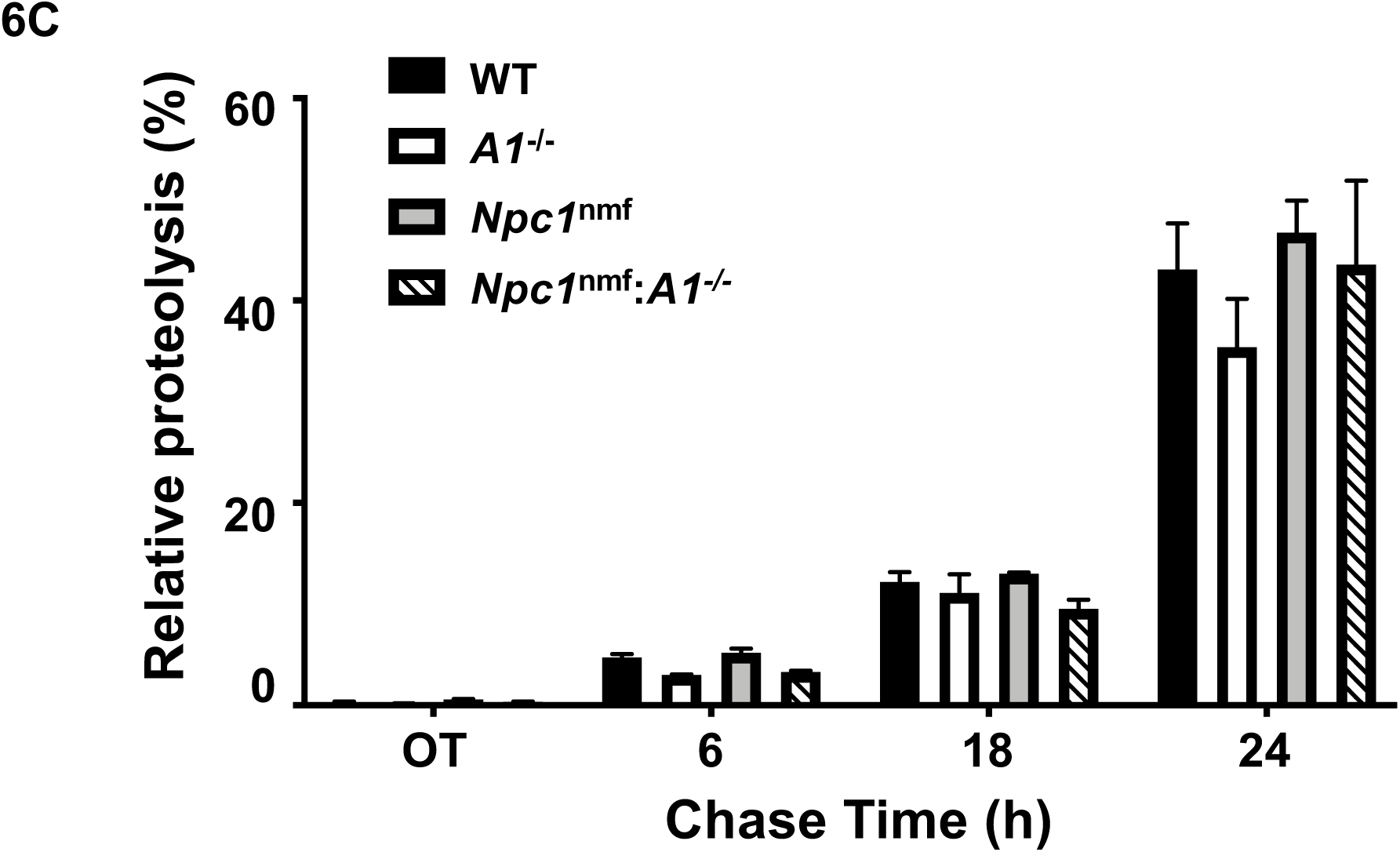
Analyses of various protein content and degradation of long-lived proteins in mouse embryonic fibroblasts (MEFs) from WT, *A1^-/-^, Npc1^nmf^*, and *Npc1^nmf^*:*A1^-/-^* mice. For parts A and B, the cell growth conditions were the same as described in Fig. 5C. Vinculin signal was used as the loading control. **A**. The relative contents of CD-M6PR protein (46-kDa) in MEFs. **B**. The relative contents of cathepsin D protein (heavy chain of the mature enzyme) (30 to 32-kDa in WT, *A1-/-*, and *Npc1nmf*:*A1-/-* cells. And 28-kDa in *Npc1nmf* cells). **C**. The degradation of long-lived proteins in MEFs from WT, *A1-/-, Npc1nmf*, and *Npc1nmf*:*A1-/-*. The analysis of protein degradation was conducted as described in the Method. Results are reported as relative proteolysis (%). Error bars indicate 1 SEM.

Cathepsin D is one of the major lysosomal enzymes that require M6PRs for processing at the TGN. From the TGN, the cathepsin D/M6PR complexes move to LE/LYS for maturation by proteolysis. Cathepsin D exists in various forms, including the precursor, intermediate, and proteolytically cleaved mature heavy chain and light chain. These forms exhibit different molecular weights on SDS-PAGE (61). Since we found that the expression level of CD-M6PR is very low in mutant *Npc1^nmf^* cells and *A1^-/-^* corrected this deficiency (**Fig. 6A**), we postulated that mutant *Npc1^nmf^* MEFs may express lower level of cathepsin D protein (the mature heavy chain), and that *A1^-/-^* in mutant *Npc1^nmf^* MEFs may correct this abnormality. To test this possibility, we performed Western blot analysis with a highly specific, monoclonal antibody against the cathepsin D mature form heavy chain. The results (**Fig. 6 B**) show that in both WT cells and *A1*^-/-^ MEF cells, this antibody recognized the mature form of cathepsin D heavy chain, which has an apparent size of 30-32 kDa. Furthermore, WT and *A1*^-/-^ cells expressed the mature form at comparable levels. In contrast, the mutant *Npc1^nmf^* cells expressed cathepsin D at a level 64% less than that of WT cells and *A1*^-/-^ cells. In addition, the size of the mature form of cathepsin D present in the *NPC1^nmf^* cells is slightly smaller (by 2-3 kDa) than that found in WT and *A1*^-/-^ cells, suggesting that within the LE/LYS of *Npc1^nmf^* cells, abnormal proteolytic cleavage of cathepsin D might occur. The *Npc1^nmf^:A1^-/-^* cells expressed cathepsin D with the same size as found in WT cells and *A1*^-/-^ cells, with protein levels higher than those found in WT and *A1*^-/-^ cells by 40%.

Together, these results show that in *Npc1^nmf^* cells, *A1*^-/-^ reversed the diminished levels of CD-M6PR and cathepsin D protein expression, consistent with both of these proteins being downstream targets of syntaxin 6 mediated vesicular trafficking. We were curious as to whether the diminished cathepsin D protein content observed in *Npc*^*nmf*^ cells may affect their ability to degrade proteins. To address this issue, we monitored the degradation of long-lived proteins by using the procedure described by Auteri *et al*. (62). The result (**Fig. 6C**) shows that, instead, the proteolysis of long-lived proteins in WT, *A1*^-/-^, *Npc1*^nmf^, and *Npc1*^nmf^ :*A1*^-/-^ MEF cells was comparable. This finding is consistent with the work of Pacheco *et al*. (63), who showed that normal and NPC1-deficient human fibroblast (Hf) cells, the degradation of long-lived proteins was comparable.

### Effects of ACAT1 inhibition on ABCA1 and NPC1 protein levels in MEFs, in mouse cerebellum, and in human fibroblasts (Hfs)

The results described above show that in *Npc1^nmf^* cells, the cathepsin D protein content is significantly decreased, and that the decrease can be corrected by *A1*^-/-^. To substantiate this finding, we sought to identify the relevant downstream targets of cathepsin D mediated signaling and decided to focus on the ATP-binding cassette transporter A1 (ABCA1). ABCA1 functions as a key cellular cholesterol efflux protein [reviewed in (64)]. It is transcriptionally regulated by liver X receptors (LXRs) (65), (66) and post-translationally regulated by various degradation mechanisms (67). In mutant *Npc1* Hfs, Choi *et al*. showed that the levels of both ACAT1 mRNA and protein are down regulated (68). In addition, in macrophages and other cells, Haidar *et al*. showed that the ABCA1 protein content is up-regulated by cathepsin D through a post-translational mechanism yet to be defined (69). In various mammalian cell lines examined, blocking ACAT1 either by genetic inactivation or by using a small molecule ACAT inhibitor increased the ABCA1 protein content; with the degree of the ACAT inhibition affecting ABCA1 levels in a cell type dependent manner (70). On the other hand, whether ACAT1 inhibition can increase ABCA1 in mutant NPC cells had not been reported previously. Since we found that mutant *Npc1^nmf^* MEF cells expressed the mature form of cathepsin D at a level significantly lower than that in WT cells (**Fig. 6B**), we postulated that this abnormality may cause mutant *Npc1^nmf^* MEF cells to express the ABCA1 protein at a lower level, and that *A1*^-/-^ may be able to correct this deficiency. To test this possibility, we performed Western blot analyses in parallel cultures of WT, *A1^-/-^, Npc1^nmf^*, and *Npc1^nmf^:A1^-/-^* MEFs. The results show that the ABCA1 protein content in mutant *Npc1* cells was lower than in WT cells or in *A1^-/-^* cells, and that *A1^-/-^* in mutant *Npc1^nmf^* cells restored ABCA1 protein content to similar level found in the WT cells (**Fig. 7A**). We prepared MEFs from WT and mutant mice that completely lacking NPC1 (the *Npc1^nih^* mouse model with a BALB/c genetic background), and found that the ABCA1 protein content in the *Npc1^nih^* MEFs was also significantly lower than that in the control WT MEF cells (**Fig. 7A; top row**; comparing the last 2 lanes on the right). These results show that in MEFs, *A1*^-/-^ restore the diminished protein content of ABCA1 in *Npc1^nmf^* cells. We also performed Western blot analyses to monitor the NPC1 protein content (**Fig. 7A; second row**) that in mutant *Npc1^mnf^* cells, NPC1 protein content was significantly lower than that in WT or *A1^-/-^* cells, confirming our previous report (46). *A1^-/-^* in mutant *Npc1* cells had a tendency to increase mutant NPC1 protein content, but the effect did not reach statistical significance (**Fig. 7A**). To serve as a control, additional results show that unlike WT MEFs, NPC1 protein was completely absent from *Npc1^nih^* MEFs (**Fig. 7A**). We next determined the level of ABCA1 mRNA in parallel cultures of WT, *A1^-/-^, Npc1^nmf^*, and *Npc1^nmf^:A1^-/-^* MEFs by RT-PCR. The results (**Fig. 7B**) show that, similar to the finding by Choi *et al*. in Hfs (68), the ABCA1 mRNA level in *Npc1^nmf^* cells is lower than that in WT cells. Furthermore, in both WT and *Npc1^nmf^* cells, *A1^-/-^* caused increase in the level of ABCA1 mRNA. This result supports the interpretation that in mutant NPC1 cells, *A1*^-/-^ restored the level of ABCA1 protein, at least in part, by restoring cathepsin D function in late endo/lysosomes.

**Fig. 7.**
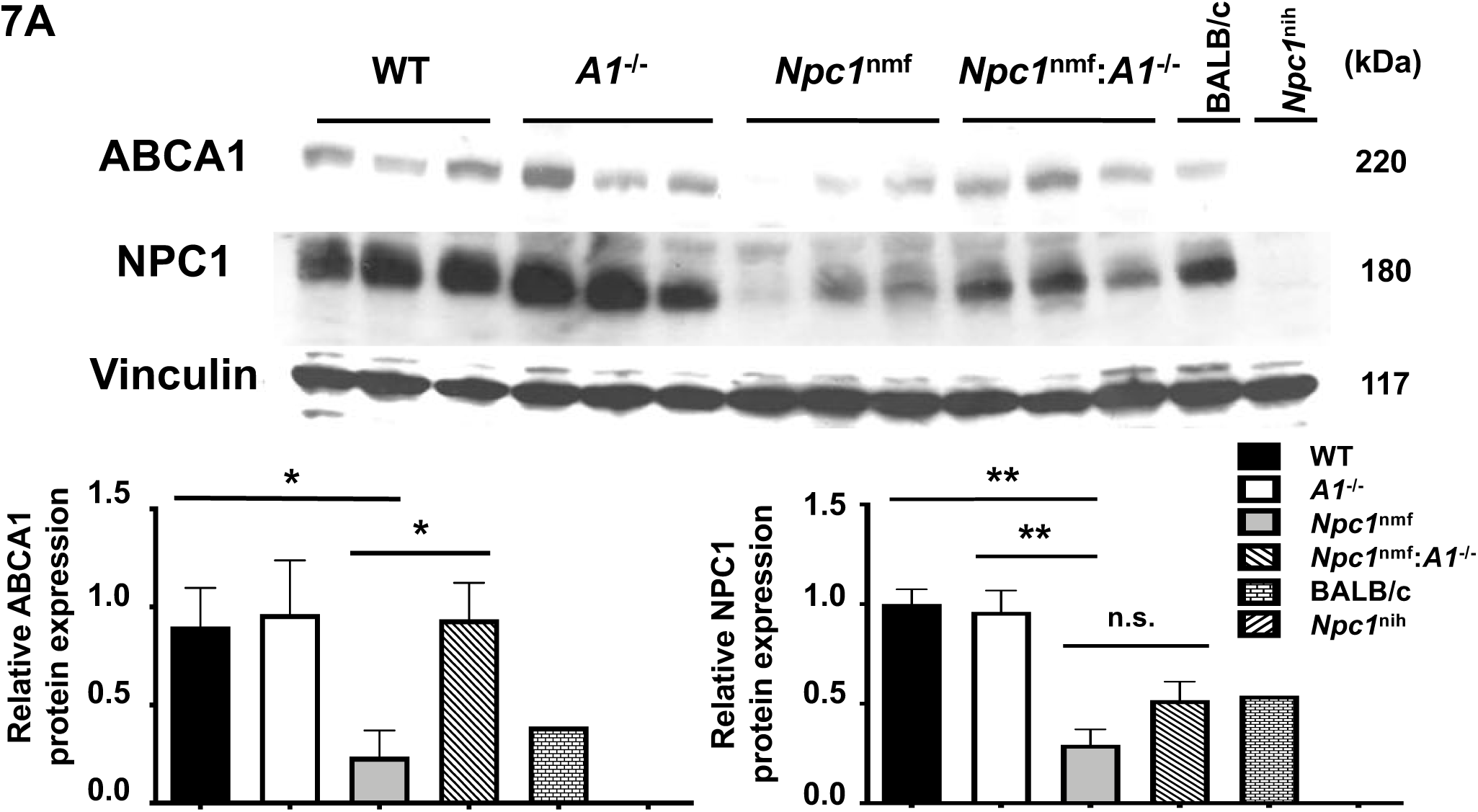
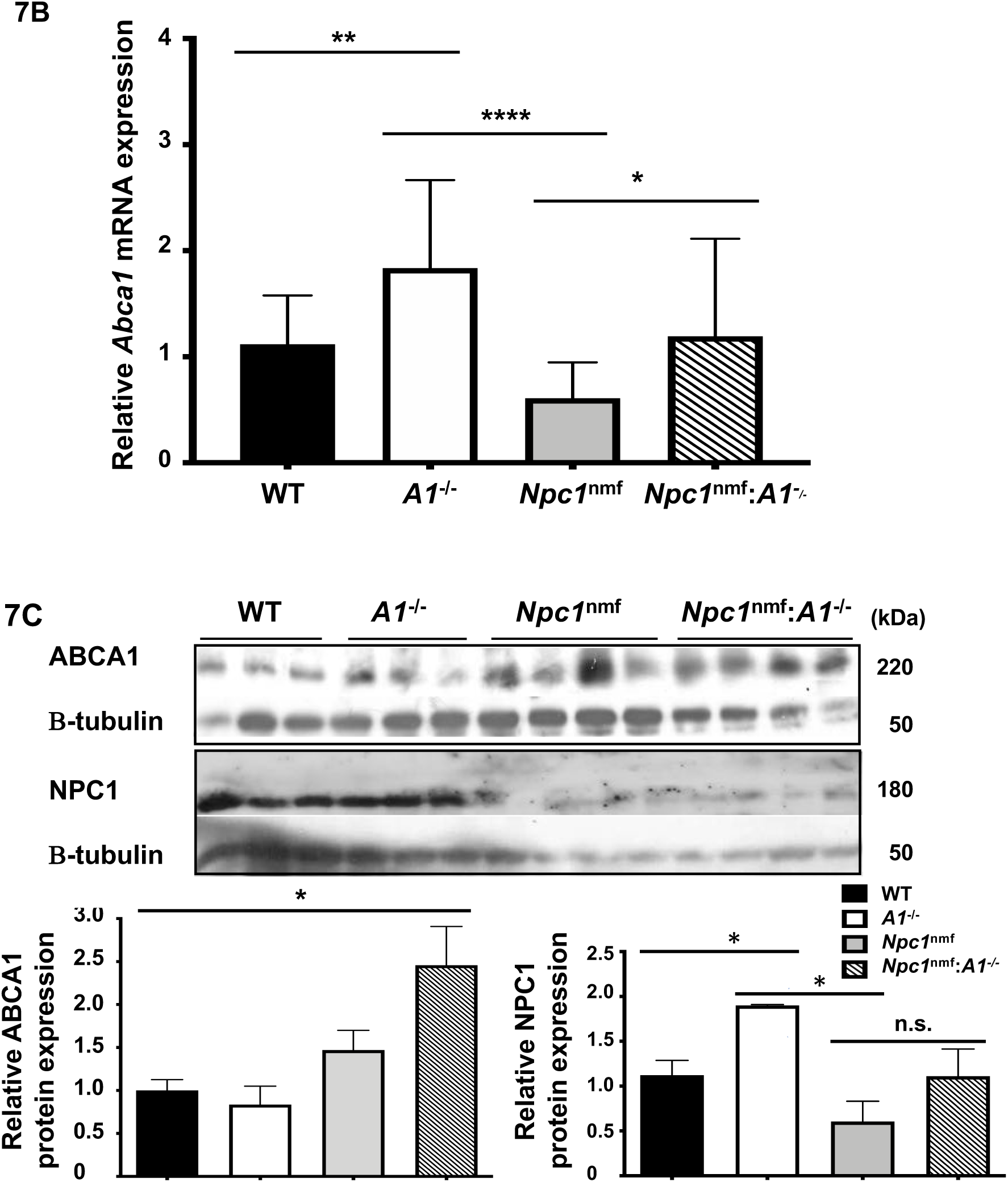
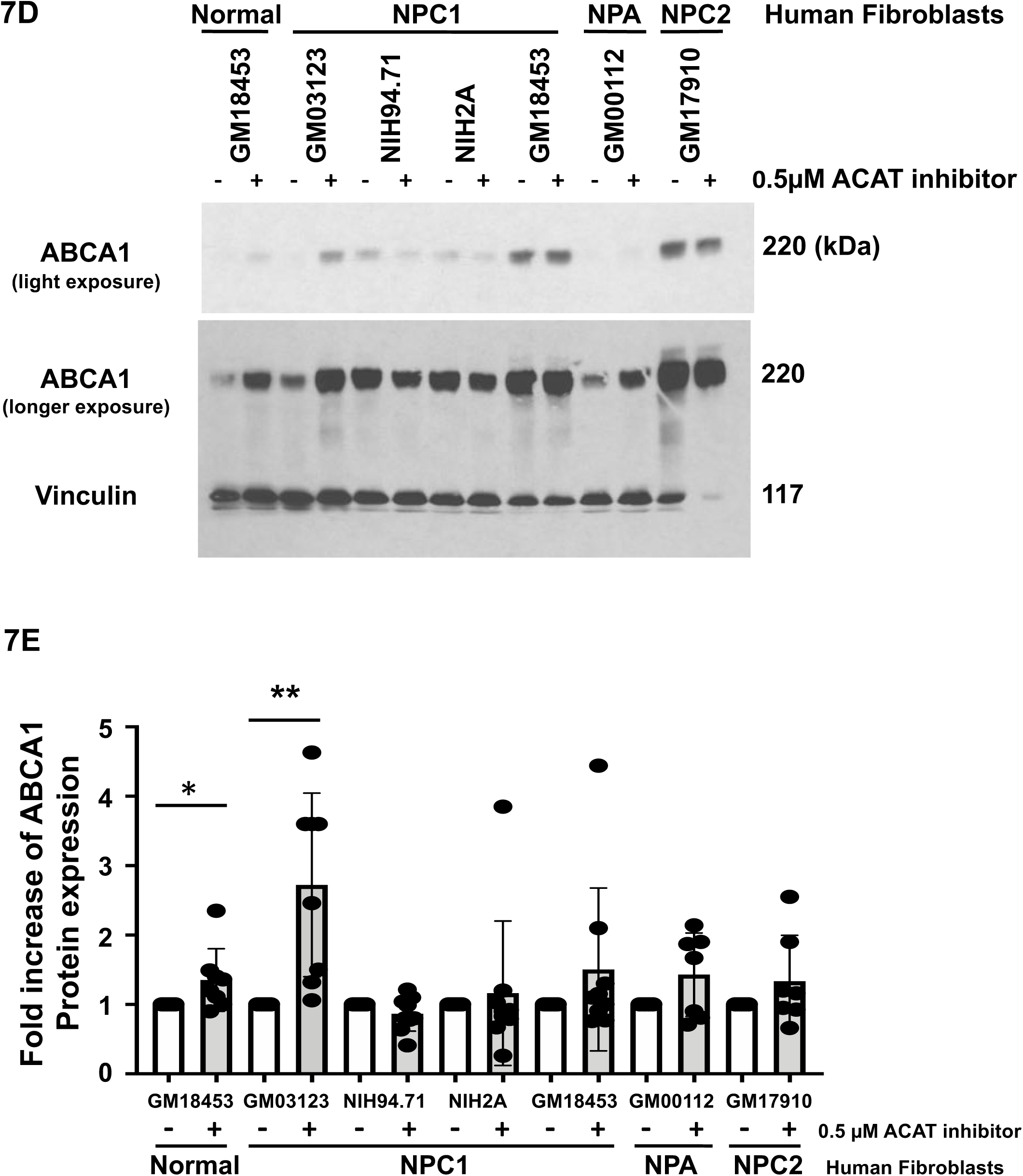
ABC1 and NPC1 expression in WT, *A1*^-/-^, *Npc1*^nmf^, and *Npc1*^nmf^:*A1*^-/-^ cells and tissues. For parts A, B, and D, the cell growth conditions were the same as described in Fig. 5C. For parts A, B, and C, cells and tissues from all four genotypes of mice (WT, *A1*^-/-^, *Npc1*^nmf^, and *Npc1*^nmf^ :*A1*^-/-^) were analyzed. **A**. Expression of ABCA1 (220 kDa) and NPC1 (180 kDa) proteins in MEFs. **B**. Expression of *Abca1* mRNA in MEFs. **C**. Expression of ABCA1 and NPC1 proteins in P80 mouse cerebellum. Tissues were prepared as described in Methods. *p*< 0.05*, n.s., not significant. **D**. Expression of ABCA1 protein in human fibroblasts (Hfs). The vinculin signal was used as the loading control. **E**. Relative expression of ABCA1 protein in human fibroblasts (Hfs) cultured in the presence or absence of 0.5 µM of ACAT inhibitor K604.

To test the *in vivo* significance of these findings, we next performed similar Western blot analyses of homogenates prepared from P80 mouse cerebellum. The results show that in mutant *Npc1* cerebellum, *A1^-/-^* significantly increased ABCA1 protein abundance (**Fig. 7C; left panel at the bottom**; comparing the 3^rd^ bar versus the 4^th^ bar). *A1^-/-^* had a tendency to also increase the mutant NPC1 protein content, but the effect did not reach statistical significance (**Fig. 7C**; right panel at the bottom; comparing the 3^rd^ bar versus the 4^th^ bar). These results demonstrate that the restorative effect of *A1^-/-^* on the low level of ABCA1 protein found in mutant NPC1 MEF can be replicated in P80 mouse cerebellum.

To determine if the beneficial effect of *A1^-/-^* on increasing the ABCA1 protein level in MEFs with an NPC mutation can also be observed in Hfs isolated from patients, we treated Hfs isolated from one normal individual, four NPC1 patients, and one Niemann-pick disease type A (NPA) patient (as indicated in **Fig. 7D**) with the small molecule ACAT1 specific inhibitor K604 at 0.5 µM for 24 h. At this concentration, K604 is expected to inhibit the ACAT1 enzyme activity in intact cells by approximately 70-80% (71). The results (**Fig. 7D, E**) show that K604 modestly increased the ABCA1 protein in normal Hf, and in one of four different mutant NPC1 Hfs (line GM03123). In the second Hf with mutant NPC1 (GM18453), in one Hf with mutant NPA (GM00112), and in one Hf with mutant NPC2 (GM17910), K604 tended to increase the ABCA1 protein, but the difference did not reach statistical significance. These results show that the enhancing effects of ACAT1 inhibition on ABCA1 protein levels observed in mutant *Npc1^nmf^* MEFs can be demonstrated in at least some Hfs with NPC1 mutations.

## Discussion

We genetically inhibited ACAT1 in a mouse model of NPC1 disease (*Npc1^nmf^*) and show that *A1*^-/-^ delays the onset of weight loss and declining sensorimotor skill, ameliorates certain systemic and neuropathological NPC disease hallmarks, and prolongs the life span by 34%. This “rescue” of *Npc1^nmf^ mice* by *A1*^-/-^ is rather surprising, and while similar attempts to extend lifespan have been made by genetic crossing of mutant NPC1 mice with mice lacking genes that encode one of the following proteins: LDL receptor (72), ApoE (73), SRBI (74), GM2 synthetase (75), GM3 synthetase (76), glucocerebrosidase 2 (77), Tau (78), and RIPK1 (a key protein that mediates necroptosis) (79), to our knowledge the current study is the first to demonstrate that knockout of a single gene can extend the life span of a mutant *Npc1* mouse model by more than 30%.

To provide a mechanistic basis for the beneficial effects of ACAT1 inhibition at the cellular level, we studied MEFs isolated from WT, *A1*^-/-^, *Npc1^nmf^*, and *Npc1^nmf^:A1^-/-^* mice, and show that in mutant NPC1 MEFs, ACAT1 inhibition alters membrane cholesterol distribution, and restores the mislocalization of syntaxin 6, a cholesterol-binding t-SNARE that normally localizes to the TGN and mediates the anterograde vesicular trafficking of the M6PRs/lysosomal hydrolase complexes. In addition to syntaxin 6, we show in mutant NPC1 cells that golgin 97, a different TGN marker, also mis-localized, and *A1*^-/-^ can restore normal golgin 97 localization at the TGN. These results suggest that mutation of NPC1 causes a certain portion of the TGN membrane to become deficient in cholesterol and that ACAT1 inhibition in mutant NPC1 cells corrects the abnormal localization patterns of both syntaxin 6 and golgin 97 by replenishing cholesterol in the TGN membrane. We also show that mutant NPC cells express diminished level of protein further downstream, such as CD-M6PR and cathepsin D (one of the key lysosomal hydrolases), and that *A1*^-/-^ restores the levels of these proteins as well. These results imply that *A1*^-/-^ restores the localization and functionality of syntaxin 6 in mutant NPC1 cells. To link the changes in cathepsin D with its downstream target(s), we also show that *A1*^-/-^ restores the diminished ABCA1 protein content observed in mutant NPC1 cells and in mutant NPC1 cerebellum. Based on these results, we propose a model to explain the actions of ACAT1 blockage: In mutant NPC cells, the inability of NPC to export cholesterol from the LE/LYS causes several membrane organelles downstream of NPC mediated cholesterol trafficking pathway to become deficient in cholesterol which leads to malfunctions in these organelles. ACAT1 resides at a certain subdomain (designated as the A1 domain) of the ER. Despite the mutation in NPC, certain cholesterol continues to arrive at the A1 domain, in an NPC independent manner, to be esterified by A1. ACAT1 inhibition causes the cholesterol pool associated with the A1 domain to translocate to other subcellular membrane compartments. The diversion of the cholesterol storage pool fulfills the needs of these membrane compartments for cholesterol and helps them regain their proper function. In the current work, we demonstrate that the TGN is one of the recipient organelles that benefits from ACAT1 inhibition. Results presented in Fig. 4A-D provide indirect evidence suggesting that in mutant NPC cells, *A1*^-/-^ may also increase the cholesterol content in other membrane compartments including the PM and perhaps the limiting membrane of the LE/LYS. Further investigations will be required to test these possibilities. It is possible that cholesterol translocation processes exist between the A1 microdomain and the microdomains present in other subcellular organelles, and that inhibition of A1 facilitates the cholesterol transfer from the A1 domain to these other membrane organelles. Multiple contact sites have been shown to exist between the ER membranes and other membrane organelles including PM, Golgi, mitochondria, and endosomes, as reviewed in (80), (81), (82), (83). The sterol transfer process between the A1 microdomain and other membrane microdomain(s) may occur through these membrane contact sites. Future work will be needed to reveal the molecular nature of the hypothetical cholesterol translocation pathways between the A1 domain and other membrane organelles.

The current study identifies ACAT1 as a new potential target for treating patients with NPC disease. It is important to evaluate the pros and cons of using small molecule ACAT inhibitors to treat NPC disease and other related diseases. At the cell culture level, studies have shown that when the cholesterol efflux process is absent (i.e., by placing cells in growth medium without any cholesterol acceptors present, or by studying cells that lack ABCA1), ACAT inhibition produced cytotoxicity (84), (33). When the cellular cholesterol efflux process is active (i.e., by including cholesterol acceptors such as apoA1 or serum lipoproteins in the growth medium), however inhibiting A1 does not cause detectable cellular toxicity (85), (86), (87). It is possible that the buildup of free (unesterified) cholesterol in cell membranes may need to reach certain threshold level before it becomes toxic to cells. At the *in vivo* level, mouse gene KO (KO) studies show that *Acat1* KO (*A1^-/-^*) mice exhibit dry skin, dry eye syndrome (88), (89), and leukocytosis (90), but their adrenal functions are normal (47) and their ability to learn and memorize are also normal (91). At the human level, at least two adult humans with presumed homozygous knockout mutations for *SOAT1* or *SOAT2* have been identified, and neither has obviously noticeable issues (92). Interestingly, recent studies have shown that, in mouse models, total *A1^-/-^* reduces pathologies associated with Alzheimer’s disease (91). Myeloid *A1^-/-^* suppresses atherosclerosis development and progression (93), (94), and suppresses diet induced obesity (95). Additional studies show that inhibiting A1 suppresses the development and progression of pancreatic cancer (96), suppresses the development of hepatocellular carcinoma (97), and potentiates the antitumor activities of cytotoxic T cells (98). Collectively, these studies suggest that if employed properly, ACAT1 may be a promising target for treating multiple human diseases. ACAT inhibitors of various structural types are available. In many cases they were developed with the original intention to treat cardiovascular diseases, and several of these inhibitors have passed a phase-I safety test for anti-atherosclerosis treatment. For safety reasons, it will be important to begin testing the ACAT inhibitors in NPC patients (and patients with other diseases) who do not also have partial deficiencies in other genes involved in the cellular cholesterol efflux process.

## Materials and Methods

### Materials

#### Chemical reagents

Fetal bovine serum was purchased from Sigma. Iron-supplemented calf serum was purchased from Atlanta Biologicals. OptiPrep was obtained from Axis-Shield. ^3^H-labeled acetate and ^3^H-labeled oleate were acquired from PerkinElmer. All chemicals (analytical grade) were purchased from Sigma-Aldrich or Fisher. Low-density lipoproteins (LDL) from fresh human blood and delipidated serum from fetal bovine serum stock were prepared as previously described (99).

#### Histological reagents

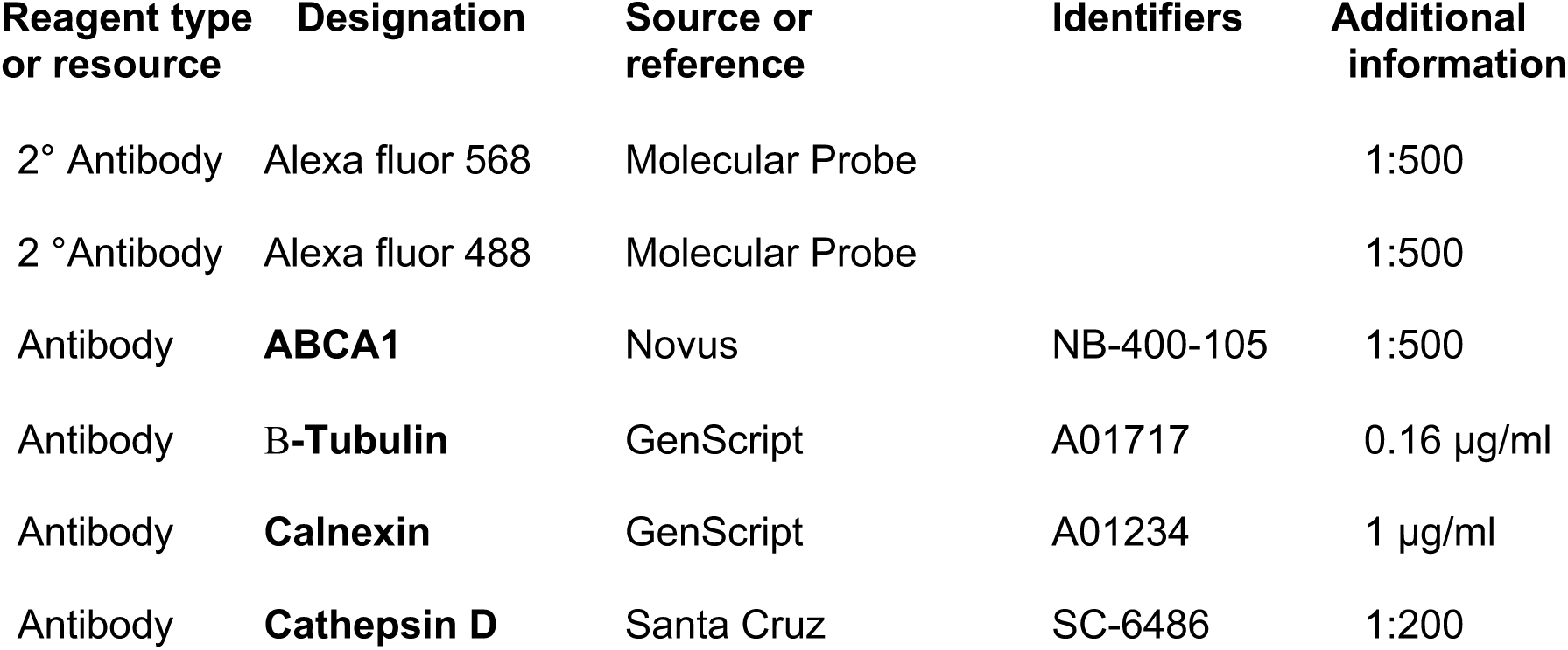
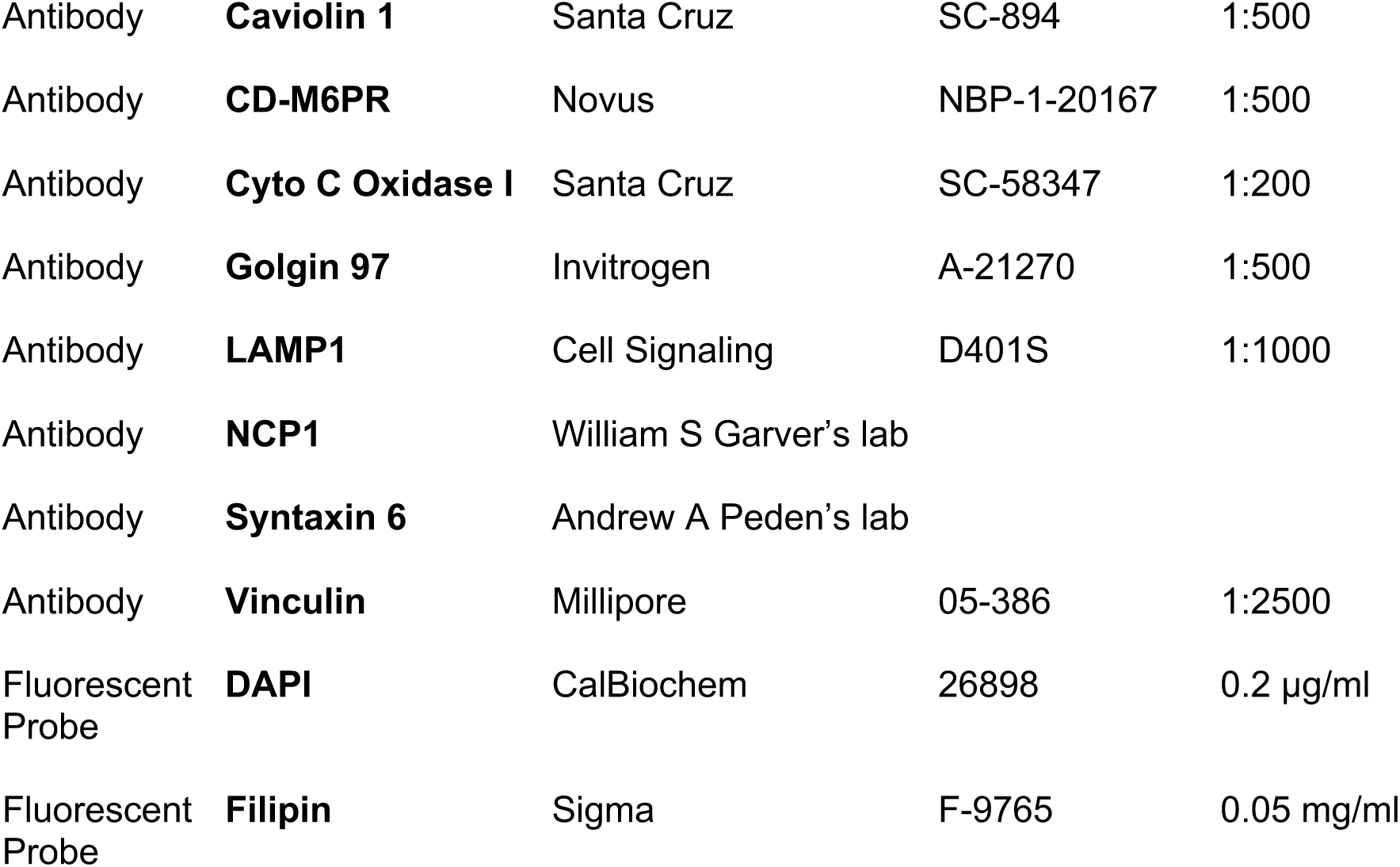

#### Primers used for ACAT1 mouse genotyping

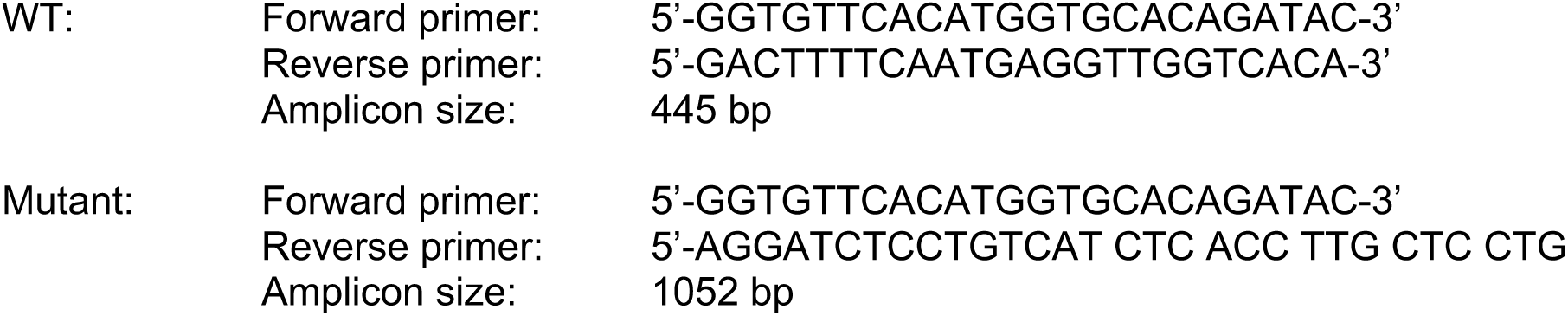

#### Primer sequences used for real time PCR (RT-PCR) analysis

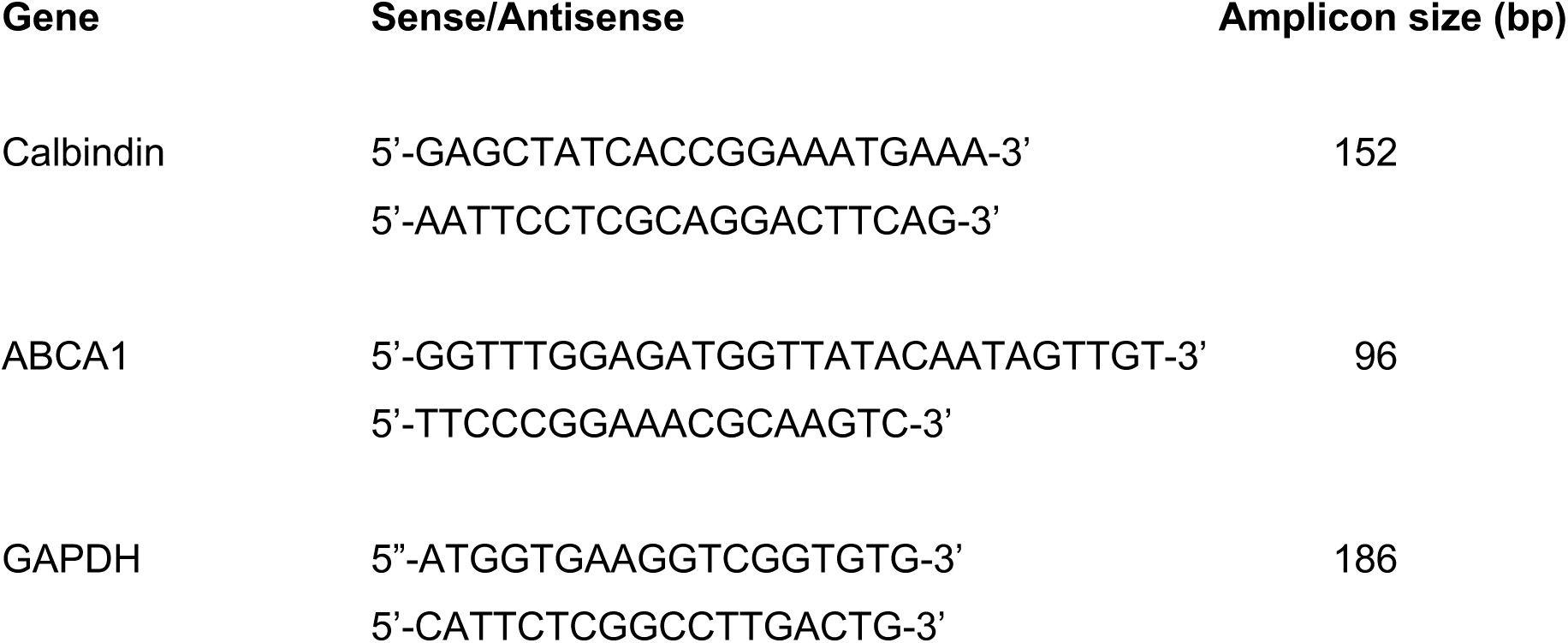

### Methods

#### Animal maintenance

Mice were fed *ad libitum* with standard chow diet, maintained in a pathogen-free environment in single-ventilated cages, and kept on a 12 h light/dark schedule, using Dartmouth Animal Research Center Institutional Animal Care and Use Committee– approved protocol number 00002020. Animals were checked daily for their entire lifespan.

When *Npc1^nmf^* mice began to have trouble reaching food, wet food pellets were placed on the bottom of their cage for the remainder of their life. Death was marked as the point where the mice could no longer ingest food or water.

#### Mouse breeding

The heterozygous mutant NPC1 mouse in C57BL/6J background (*Npc1^nmf/wt^*:*Acat1^+/+^*mouse; from Jackson laboratories) was crossed with *Acat1*^-/-^ mouse (global ACAT1 KO mouse) [(47); received from Dr. Sergio Fazio in C57BL/6J background], to produce *Npc1^nmf/wt^*:*Acat1*^-/-^ mice and *Npc1^nmf/wt^*:*Acat1*^+/+^mice. After two rounds of breeding, the resultant *Npc1^nmf/wt^*:*Acat1^-/-^* mice were set up as breeding pairs to generate the *Npc1^nmf/nmf^*:*Acat1^-/-^*mice (Designated as *Npc1^nmf^* :*A1^-/-^* mice) and *Npc1^wt/wt^*:*Acat1^-/-^* mice (Designated as *A1^-/-^* mice). The *Npc1^nmf/wt^*:*A1^+/+^* mice were set up as breeding pairs to generate *Npc1^nmf/nmf^*:*A1^+/+^*mice (Designated as *Npc1^nmf^* mice) and *Npc1^wt/wt^*:*A1^+/+^*mice (Designated as WT mice).

#### NPC1nmf mouse genotyping

The protocol described by (46) was followed with minor modification: 10 µl of reaction buffer containing 10 ng of mouse-tail genomic DNA, 1x TaqMan genotyping master mix, and 1x SNP Custom TaqMan SNP assay mixture. The PCR reaction was carried out by amplifying at 95°C for 5 min, followed by 45 cycles of: 92°C for 15 s and 60°C for 1 min.

#### ACAT1 mouse genotyping

PCR conditions were: 94°C for 1.5 min, followed by 35 rounds of: 94°C for 30 s, 62°C for 60 s, 72°C for 60 s. Lastly, 72°C for 2 min. The primers used are described above in the Materials.

#### Mouse motor skills

Mouse motor skills were assessed by RotaRod test using a commercially available instrument (purchased from Med Associates Inc, Fairfax, VT) in a manner similar to what was previously reported (100), with slight modifications. Briefly, after a brief initial training period, mouse motor skills were monitored from six weeks postnatal age until failure. Each week mice were given three consecutive trials on a constant speed rotarod at 24 rotations per min for up to 90 s for each of the three trials. WT and *A1^-/-^* mice passed all trials running for at least 10 s on any of the three consecutive trials during every week assessed. Age of rotarod test failure in *Npc1^nmf^* and *Npc1^nmf^:A1^-/-^* mice was measured as the age at which mice failed to run on the rotarod for at least 10 s during at least one of the three consecutive trials.

Rotarod trial failure included falling off the rod before 10 s or freezing and clasping to the rotarod and not running or moving.

#### Histological analyses

Hematoxylin and Eosin staining of mouse liver, spleen and lung tissues, and Purkinje neurons in cerebellum at postnatal day 80 were performed by the Histology Service at the Jackson Laboratory, using standard protocols in a Leica Autostainer XL automated processor.

#### Cell culture

MEFs were isolated according the procedure described (101). MEF were grown as monolayers at 37°C with 5% CO_2_ in DMEM supplemented with 10% serum and MEM non-essential amino acids (Gibco), or with 5% delipidated fetal bovine serum and 35 μM oleic acid, and with penicillin/streptomycin. Each experiment was performed with cells grown in triplicate dishes.

#### Degradation of long-lived proteins in MEFs

The procedure described in (62), (63) was adopted with minor modification. MEFs were seeded on 12-well plates at a density of 0.012×10^6^ cells in triplicate. Media were replaced the night before the experiment. Cells were rinsed with 2 ml of MEM and labeled with 1 ml of 2 µCi/ml of ^3^H leucine in MEM+10% serum. At time zero, cells were washed twice with HBSS, then chased by 1 ml MEM+2.8 mM leucine without serum. At each time point indicated, media were transferred to a microcentrifuge tube, trichloroacetic acid added to a final concentration of 20% and BSA to a final concentration of 3 mg/ml.

Samples were incubated at 4°C for 1 h, centrifuged at 15,000g at 4°C for 5 min, supernatants and pellets were collected. Ecoscint H was added for scintillation counting. Cells were washed with PBS and incubated in 0.1 M NaOH in 0.1% deoxycholate for 1 h. 40 µl in duplicates were aliquoted for protein determination.

#### Western blot analyses

From either freshly isolated mouse cerebellum tissues or tissue culture cells: it was prepared in either 10% SDS (for syntaxin 6, CD-M6PR, NPC1, or cathepsin D), or RIPA buffer (for ABCA1), plus protease inhibitor (Sigma), and homogenized in a stainless-bed Bullet Blender twice for 3 min each at 4°C. Homogenized lysates were run on a 6% gel (for ABCA1), 10% gel (NPC1, LAMP1) or 12% gel (for Syntaxin 6, CD-M6PR, Cathepsin D, Caveolin 1, and Cytochrome C Oxidase), and transferred to PVDF membrane in Towbin buffer. Signal intensities were normalized to vinculin (117-kDa) or **B** tubulin (50-kDa) expression by NIH Imaging software.

#### RNA isolation and real-time PCR (RT-PCR) experiments

Total RNA was isolated from TRIzol reagent (Invitrogen) following the manufacturer’s instruction. The RNA was dissolved in sterile water treated with DNaseI (Ambion). 1 µg of RNA was used to synthesize cDNA, according to the instructions in the BioRad iScript cDNA Synthesis Kit. Real-time PCR was carried out using the iTaq Universal SYBR Green Supermix (from Bio-Rad) with the Applied Biosystems Step One RT-PCR system. Relative quantification was determined by using the delta CT method. The primer sequences used are listed above in the Materials. The PCR reaction conditions were as described previously (91).

#### Subcellular fractionation

Procedures were carried out as described previously (22). Cells grown in one 150-mm dish to near confluence were washed twice with phosphate-buffered saline, once with homogenization buffer (HB, 250 mM sucrose, 20 mM Tris-HCl, pH 7.4, 1 mM EDTA), harvested to 1 ml HB with protease inhibitors, and homogenized by using a stainless-steel homogenizer with 40 strokes. The post-nuclear supernatants were placed onto the top of an 11 ml 5-25% OptiPrep discontinuous gradient in HB. The gradient was centrifuged at 200,000 x g (40,000 rpm) for 3 h in a Beckman SW41 rotor; 15 fractions (800 µl each) were collected from the top.

#### Lipid syntheses in intact cells

The analysis of cholesterol biosynthesis in intact cells was carried out as previously described (102), exposing cells to ^3^H-labeled acetate for 1 h followed by lipid extraction. The measurement of cholesterol esterification in intact cells was performed by exposing cells to ^3^H-labeled oleate/BSA for 1 h followed by lipid extraction and analysis, as previously described (103).

#### Fluorescence microscopy

MEF cells were cultured on poly-D-Lysine (70-150kDa) glass coverslips (MatTek) in 12-well plates for 24 h, and fixed in 4% paraformaldehyde (EMS) at RT for 10 min. After washing with PBS, the cells were permeabilized with 0.3% Triton X-100 for 20 min. Cells were then washed with PBS prior to blocking with 5% goat serum in PBS for 1 h at RT, followed by staining with primary antibodies for at least 1 h at RT. Cells were washed with PBS before and after incubation with Alexa Fluor 568 or Alexa Fluor 488 as the secondary antibodies at 1:500 dilution for 1 h at RT, DNA was counter stained with DAPI (0.2 µg/ml) at RT for 10 min to visualize the nucleus.

To identify cellular cholesterol, cells were fixed in 4% paraformaldehyde (without the use of detergent), after several washes, cells were pre-incubated with 1.5 mg/ml glycine in PBS for 10 min at RT, then incubated with Filipin (50 µg/ml) at RT for 1 h as previously described (104).

Images were acquired by using the Andor W1 Spinning Disk Confocal system (Nikon Eclipse Ti inverted microscope, and Andor Zyla camera), with a 60x oil-immersion lens, using three laser lines (403-nm laser for DAPI, 488-nm and 561-nm filters for FITC and Texas Red respectively). Z-stacked fluorescent images were taken by 11 optical slices at 0.2 µm intervals to enhance the spatial signal allocation. Images were visualized by using Fiji-Image J Software, and processed using Nikon Elements to create the “Maximum Intensity Projection”, and calculate the Pearson’s correlation coefficient. The Reflex angle was determined according to Mitchel *et al*. (55), as the angle subtended by the edges of the positive fluorescence signals, using the center of the nucleus (based on the DAPI positive signal) as the vertex.

#### Statistical analysis

Statistical comparisons were made by using a two-tailed, unpaired student *t* test according to GraphPad Prism 8. Difference were considered significant when the *P* value was less than 0.05. (*p* < 0.0001****, *p* < 0.001 ***, *p* < 0.01 **, *p* < 0.05 *). n.s. indicates the differences were not significant.

## Author contributions and Conflict of interest statement

M.A.R., C.C.Y.C., and T.-Y.C. designed research; C.C.Y.C., M.A.R., E.M.M., and M.H., P.S performed research; R.A.M., A.P., and W.G. contributed new reagents/analytic tools; C.C.Y.C., M.A.R., R.A.M. and T.-Y.C. analyzed data; T.-Y.C., M.A.R., and C.C.Y.C., wrote the paper. All authors approved the final version of the manuscript.

The authors declare no conflict of interest.

## Acknowledgments

We thank Drs. Ann Lavanway, Wen-Lih Lee, Safia Omer, Zdenek Svindrych and Lona Young for advice in confocal microscopy usage, we thank Dr. Henry Higgs for providing the anti-golgin 97 antibodies. We thank Yohei Shibuya and other members of the Chang lab for helpful discussions during the course of this work. We also thank Dr. Gustav Lienhard for careful reading of this manuscript.

This work was supported by NIH grant R01 AG063544 (to T.Y. Chang and C.C.Y. Chang). We acknowledge the shared facilities of the pre-clinical imaging and microscopy resource at the Norris Cotton Cancer Center at Dartmouth with NCI Cancer Center Support Grant 5P30 CA023108-41, and a grant P20-GM113132 to support Institute for Biomolecular Targeting at Dartmouth. The content is solely the responsibility of the authors and does not necessarily represent the official views of the National Institutes of Health.

## Abbreviations

ABCA1: ATP-binding cassette protein A1
ACAT/SOAT: Acyl-coezyme A:cholesterol acyltransferase/sterol O-acyltransferase
KO: gene knockout;
*A1*^-/-^, *Acat1*^-/-^: ACAT1 gene ablation
CD-M6PR: cation-dependent mannose-6-phosphate receptors
ER: endoplasmic reticulum
PM: plasma membrane
TGN: trans-Golgi network;
LE: late endosomes
LXRs: liver X receptors
NPC: Niemann-Pick type C
*Npc1^nmf/nmf^, Npc1^nmf^, or Npc1^nmf164^*: An *Npc1* disease mouse model with D1005G mutation
NPA: Niemann-Pick type A.

